# A conserved small RNA-generating gene cluster undergoes sequence diversification and contributes to plant immunity

**DOI:** 10.1101/2025.07.20.665670

**Authors:** Li Feng, Yingnan Hou, AmirAli Toghani, Zhixue Wang, Bozeng Tang, Nicola Atkinson, Hui Li, Yue Qiao, Yan Wang, Jian Hua, Jixian Zhai, Wenbo Ma

## Abstract

Small RNA-mediated gene silencing contributes to plant immunity. The secondary small interfering RNA (siRNA) pathway promotes defense by silencing target genes in invading fungal and oomycete pathogens. Many secondary siRNAs derive from transcripts potentially encoding pentatricopeptide repeat (PPR) proteins. Here, we report that siRNA production is an ancient function of an evolutionarily conserved clade of *PPR* genes that undergo extensive within-species diversification. In *Arabidopsis thaliana*, siRNA-source *PPR*s are physically clustered in one locus on Chromosome 1. These sequences are diversified through gene duplication followed by sequence diversification as well as accumulation of high-impact variations including pseudogenization. This diversity leads to the accumulation of a diverse *PPR*-siRNA pool, consistent with an engagement in a co-evolutionary arms race with the pathogens. This study defines siRNA-producing PPRs as a family of defense genes and highlights the potential of *PPR*-siRNA-based engineering for enhancing broad-spectrum disease resistance.

## Main

Plants employ a sophisticated immune system to combat pathogens. Pathogen perception activates immune signalling that leads to various defense responses such as cell wall reinforcement, reactive oxygen species (ROS) production, and release of antimicrobial molecules ^1^. Recent studies suggest that small RNAs (sRNAs) can function as antimicrobial agents by directly targeting pathogen genes through a process termed host-induced gene silencing (HIGS) ^2^.

sRNAs are short, non-coding RNA molecules that repress target gene expression through nucleotide base pairing ^3^. Plant sRNAs are classified into two major categories, microRNAs (miRNAs) and small-interfering RNAs (siRNAs) ^4^. miRNAs originate from *MIRNA* (*MIR*) genes, whose primary transcripts fold into imperfectly paired stem-loop structures ^5^. siRNAs, on the other hand, arise from perfectly matched double-stranded RNAs (dsRNAs) generated by RNA-dependent RNA polymerases (RDRs) ^6^. Dicer-like (DCL) proteins process *MIR* stem-loops or RDR-derived dsRNA precursors into miRNA or siRNA duplexes, which are subsequently loaded into ARGONAUTE (AGO) protein complexes for target gene silencing. Some siRNAs require cleavage of specific transcripts guided by miRNAs to initiate their biogenesis and are thereby named secondary siRNAs. The cleaved RNAs are then used as templates to produce dsRNA precursors, which are further processed into an array of 21 or 24 nt siRNA duplexes ^7^. These “secondary” siRNAs are spaced in phased intervals and are therefore also called phased siRNAs or phasiRNAs. Sequences that give rise to phasiRNAs include the non-coding *TAS* genes and protein-coding genes, especially those predicted to encode Nucleotide-binding leucine-rich repeat domain (NLR) and pentatricopeptide repeat (PPR) proteins ^8^.

The functional significance of the secondary siRNA pathway lies in the potential widespread gene silencing through a cascade in which secondary siRNAs could target genes not regulated by the primary miRNAs, hence amplifying the silencing efficacy. This is believed to be particularly effective in regulating large gene families, such as NLRs and PPRs ^7^. In particular, NLRs are canonical immune receptor proteins and siRNAs produced through the miRNA-*NLR*-siRNA and/or miRNA-*TAS*-*NLR*-siRNA pathways potentially have a global impact on *NLR* gene expression ^8–11^. For example, in soybean, potato, and barley, members of the miR482/miR2188 superfamily trigger production of secondary siRNAs that presumably potentially silence a large number of *NLR* genes to prevent fitness-reducing autoimmunity in the absence of pathogen attack ^12^. Reduced accumulation of miR482/miR2118b in tomato led to enhanced resistance to the oomycete pathogen *Phytophthora infestans* and the bacterial pathogen *Pseudomonas syringae*, likely due to decreased production of *NLR*-silencing siRNAs and subsequently, increased expression of NLRs ^10^. More recently, *trans*-acting siRNAs (tasiRNAs) generated from *TAS3* were found to regulate systemically acquired resistance by modulating the expression of auxin response factors ^13^.

In addition to regulating endogenous gene expression, secondary siRNAs have been found to elevate defense by silencing genes in invading eukaryotic pathogens through HIGS ^2^. In *Arabidopsis thaliana*, the accumulation of siRNAs derived from *PPR* transcripts increases during the infection of the oomycete pathogen *Phytophthora capsici* ^14^. Mutations in miR161, which targets a small subset of *PPR* transcripts and triggers siRNA production, lead to hypersusceptibility to *P. capsici*, suggesting a contribution of *PPR*-derived siRNAs (*PPR*-siRNAs) to defense. *A. thaliana PPR*-siRNAs represent a diverse pool with thousands of sequences that are predicted to target many *P. capsici* genes. Silencing of one of the predicted targets was confirmed during infection ^14^. These findings support a role of *PPR*-siRNAs in plant defense. Consistent with this, oomycete pathogens encode silencing suppressors that contribute to virulence ^14,15^.

*PPR*s constitute a large gene family in plants, with over 400 members in most sequenced species ^16^. Functionally characterized PPR proteins have sequence-specific nucleotide-binding activity and act as molecular adaptors that direct RNA processing complexes to specific transcripts in mitochondria and chloroplast ^17^. A subset of *PPR* genes can be recruited to the miRNA-triggered secondary siRNA pathway and this phenomenon seems to be prevalent in eudicots ^18^. The function of *PPR*-siRNAs in plant defense suggests that the siRNA-source loci should be engaged in a co-evolutionary arms race with the pathogens, thus may exhibit a distinct evolutionary trajectory ^19^. Interestingly, most siRNA-producing genes in the *A. thaliana* accession Col-0 cluster together and distantly from other *PPR* genes in a phylogenetic tree generated from ∼300 PPR proteins ^20^. Whether this pattern is common in *A. thaliana* and other plants, and how they are correlated with siRNA production, is unknown.

In this study, we systematically analyzed siRNA-producing *PPR* genes in eight *A. thaliana* accessions and expanded this analysis to a wide range of evolutionarily diverse plant species. In *A. thaliana*, we found that siRNA-producing *PPR*s are physically and phylogenetically clustered. The siRNA-producing *PPR*s, but not those do not give rise to siRNAs, exhibit a high level of sequence diversity, consistent with their engagement in host-pathogen co-evolution. Despite the sequence variation, siRNA-producing *PPR*s retain the target sites of miR161 and several tasiRNAs that can trigger secondary siRNA production, resulting in a diverse pool of *PPR*-siRNAs. Functional analysis supports the hypothesis that siRNA production, not putative protein products, is responsible for the disease resistance function of a siRNA-source *PPR* gene. Importantly, PPR analysis of evolutionarily diverse plant species revealed a dedicated “siRNA-producing” clade that exhibits within-species diversification, suggesting that siRNA production is an ancient function of a specific phylogenetic group of *PPR* genes. Finally, we show that heterologous expression of an *A. thaliana PPR*-siRNA-producing cassette can enhance *Phytophthora* resistance in *Nicotiana benthamiana*. This study defines siRNA-producing *PPR*s as a new family of defense genes and provides a proof-of-concept strategy to elevate disease resistance by engineering *PPR*-siRNAs.

## Results

### A dedicated siRNA-producing *PPR* gene cluster in *A. thaliana*

We investigated *PPR*-siRNA populations and siRNA-producing *PPR* loci in eight *A. thaliana* accessions with high-quality genome assemblies, including An-1, Col-0, Ct-1, Cvi-1, Eri-1, Kyo-1, L*er*-0, and Sha ^21–23^, using sRNA-seq. After quality control and exclusion of structural RNA-derived reads, we obtained 1 to 7 million genome-matched reads from each accession in the size range of 18-28 nt, comprising 0.4 to 1 million unique reads (**Table S1**). Size distribution of sRNA population in each accession exhibited two prominent peaks at 21-nt and 24-nt (**Fig. S1a**), consistent with typical patterns observed in *A. thaliana* ^24^.

Using this dataset, we determined *PPR* loci from which siRNAs were spawned. For this purpose, PPR-encoding genes were identified from each *A. thaliana* accession based on sequence similarity with a set of 475 *PPR* genes defined in Col-0 ^25^. Using SimpleSynteny ^26^, blastn, and blastp ^27^, 467, 471, 459, 465, 464, 467, and 461 *PPR* genes were annotated in An-1, Ct-1, Cvi-1, Eri-1, Kyo-1, L*er*-0, and Sha, respectively (**Table S2**). Total siRNAs extracted from these *PPR* loci showed a predominant peak at 21-nt in all accessions (**Fig. S1b**), consistent with the previous observation in Col-0 ^20^.

To identify siRNA-producing *PPR*s, we calculated the normalized accumulation (transcripts per million, TPM) of siRNAs derived from each *PPR* gene in these *A. thaliana* accessions. Using a cutoff of TPM > 5, 127 *PPR* genes were identified as siRNA-producing loci (**Fig. 1a**; **Table S3**). While the numbers of non-siRNA-producing *PPR*s remain relatively constant in all accessions (from 448 to 455, with a standard deviation of 2.33), the numbers of siRNA-producing genes are more variable (from 10 to 22, with a standard deviation of 3.52). Furthermore, for non-siRNA-producing *PPR*s, one gene in Col-0 has a single homolog in each of the other seven accessions in almost all cases. In contrast, siRNA-producing *PPR*s often have multiple homologous genes in another accession (**Fig. S1c**; **Table S3**). These observations indicate an increased variability in siRNA-producing *PPR*s among the accessions.

**Figure 1.**
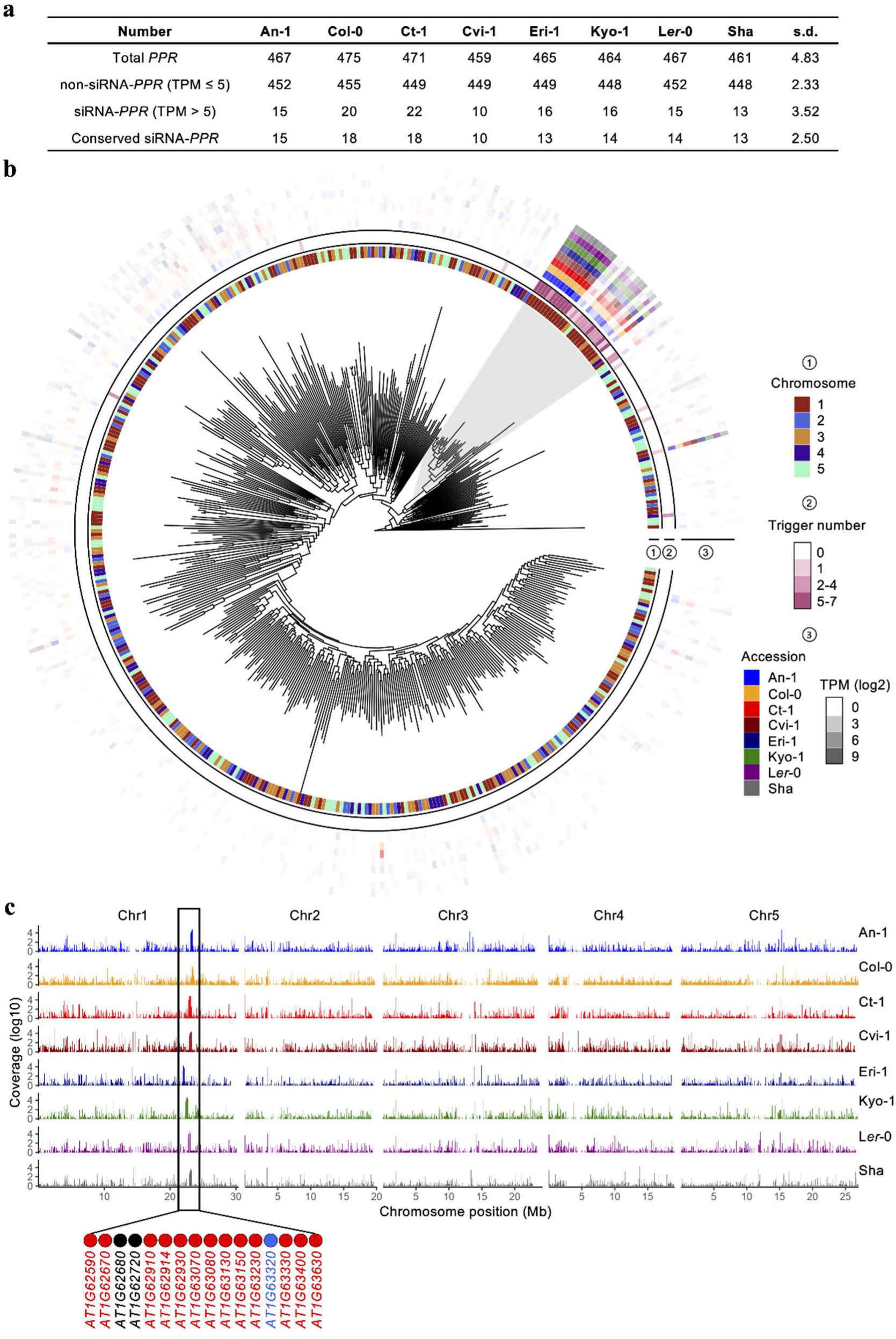
A designated *PPR* gene cluster produces siRNAs in *A. thaliana*. **a.** siRNA-producing *PPR* (siRNA-*PPR*) homologs identified in eight *A. thaliana* accessions. Conserved siRNA-*PPR*s were defined as *PPRs* that accumulated siRNAs with a TPM (Transcripts Per Million) value > 5 in at least five accessions. s.d. represents the value of standard deviation. **b.** Phylogenetic analysis of the complete set of 475 PPRs in *A. thaliana* Col-0 using amino acid sequences. Ring 1 indicates the chromosomal location of each PPR-encoding gene. Ring 2 indicates the number of sRNA triggers predicted from each *PPR* transcripts. These sRNA triggers include miR161.1, miR161.2, miR400, TAS1a 3′ D9 (-), TAS1b 3′ D4 (-), TAS1c 3′ D6 (-), TAS1c 3′ D10 (-), TAS2 3′ D6 (-), TAS2 3′ D9 (-), TAS2 3′ D11 (-), and TAS2 3′ D12 (-). Ring 3 presents the abundance of 21-nt siRNAs derived from each *PPR* homologs in the eight *A. thaliana* accessions. siRNA abundances are represented by log_2_-transformed normalized TPM. Highlighted in grey is one phylogenetic group that produces the majority of *PPR*-siRNAs. **c.** The majority of *PPR*-siRNAs are produced from a conserved gene cluster on Chromosome 1. The entire complement of 21-nt siRNAs derived from coding sequences in each of the eight *A. thaliana* accessions are mapped to their chromosomal locations. A major peak is highlighted on the longer arm of chromosome 1, representing a conserved sRNA-producing hotspot that harbors multiple siRNA-producing *PPR* genes. Red dots label siRNA-producing *PPR*s that belong to the same phylogenetic group highlighted in panel **b**; the gene with a blue dot produces siRNAs but is not clustered within the phylogenetic group highlighted in panel **b**; the black dots label non-siRNA-producing *PPRs*.

Twenty *PPR*s produced siRNAs in Col-0. Mapping these siRNA-producing *PPR*s on a phylogenetic tree generated using the amino acid sequences of the entire set of 475 PPRs revealed that 18 of the siRNA-producing *PPR*s belong to the same clade (**Fig. 1b**). This result is similar to the previous analysis using a smaller set of PPRs ^19^. These 18 genes are responsible for the production of 97.95% of the total 21-nt *PPR*-siRNA population in Col-0.

The initial step of secondary siRNA production involves targeting of siRNA-producing transcripts by trigger sRNAs, being miRNA or siRNAs, via sequence complementarity ^28,29^. Eleven miRNAs and tasiRNAs have previously been identified as potential triggers of *PPR*-siRNA production in *A. thaliana* through the miRNA-*PPR*-siRNA or miRNA-*TAS*-*PPR*-siRNA pathways ^18,20^. These trigger sRNAs include miR161.1, miR161.2, miR400, and eight tasiRNAs derived from *TAS1* and *TAS2*. Target site prediction suggests that the 18 siRNA-producing *PPR*s belonging to the same phylogenetic group are highly enriched with target sites of the 11 sRNA triggers (**Fig. 1b**; Ring 2). Many gene members are predicted to contain target sites of more than five trigger sRNAs, correlated with their role as the main *PPR*-siRNA producers in the genome.

We next determined the conserved siRNA-producing *PPR*s using a criterium that siRNA production must be detectable in at least five *A. thaliana* accessions from the same homologous gene. This analysis results in the identification of 115 genes in the eight accessions that we named as “core” siRNA-producing *PPR*s (**Fig. 1a**). Seventeen of the 18 core siRNA producers in Col-0 belong to the same phylogenetic group highlighted in **Fig. 1b**. Remarkably, homologs of these 17 genes in the other accessions are also the most prominent siRNA producers in their specific accession, suggesting that siRNA-production is a conserved feature of this phylogenetic group in *A. thaliana* (**Fig. 1b**; Ring 3).

We noticed that all siRNA-producing *PPRs* in this phylogenetic group, except for one, are located on Chromosome 1 (**Fig. 1b**; magenta-colored grids in Ring 1). To further investigate how these genes may be physically linked, we conducted a genome-wide analysis and mapped the entire population of 21-nt siRNAs derived from coding sequences in each *A. thaliana* accessions. The results revealed a prominent peak on the long arm of Chromosome 1 as a major siRNA-producing hotspot in a similar location of all accessions (**Fig. 1c**). In Col-0, 16 *PPR* genes are located within a region of 0.4 Mb (nucleotides 23,179,248 – 23,587,005), among which include 14 siRNA-producing *PPR*s (**Fig. 1c**). All of these 14 genes, except for one (AT1G63320), belong to the same phylogenetic group highlighted in **Fig. 1b**. siRNAs produced from this gene cluster account for 78.76% to 98.33% of the total *PPR*-siRNAs across the eight *A. thaliana* accessions (**Fig. S1d**). Taken together, these analyses identified a phylogenetically related and physically linked *PPR* gene cluster that serves as a conserved siRNA-producing hotspot in *A. thaliana*.

### Genetic dynamics in the siRNA-producing *PPR* cluster

Physical clustering of genes belonging to the same family may accelerate their evolution. For example, genes encoding immune receptors, especially the *NLR* genes, often form clusters in the genome ^30^. Some *NLR* gene clusters potentially undergo asymmetrical expansion, with certain members exhibiting a greater tendency for duplication and diversification than others ^31^. We examined potential gene duplication events among 87 siRNA-producing *PPR* genes located in the siRNA-producing *PPR* gene cluster on Chromosome 1 (called “hotspot” siRNA-producing *PPR*s hereafter) by assessing their sequence similarity across the eight *A. thaliana* accessions. In addition, homologs of the siRNA-producing *PPR AT5G16640* of Col-0, which is located on Chromosome 5 but phylogenetically clustered with the hotspot siRNA-producing *PPR*s, were included in the analysis for comparison.

The hotspot siRNA-producing *PPR*s can be categorized into two groups (using a cutoff of > 75% nucleotide sequence similarity) that are also clearly separated from the *AT5G16640* homologs (**Fig. 2a**). The largest group (G1) consists of 75 genes as homologs of 12 hotspot siRNA-producing *PPR*s in Col-0. G2 consists of 12 hotspot siRNA-producing *PPR*s that are homologous to the Col-0 genes *AT1G63230* and *AT1G63630*. Although variable numbers of genes in different accessions belong to G1 and G2, G3 is comprised of eight homologs of the Col-0 *AT5G16640*, one in each accession (**Fig. 2a**). The gene number variation in different accessions indicates a higher degree of dynamics among the hotspot siRNA-producing *PPR*s.

**Figure. 2.**
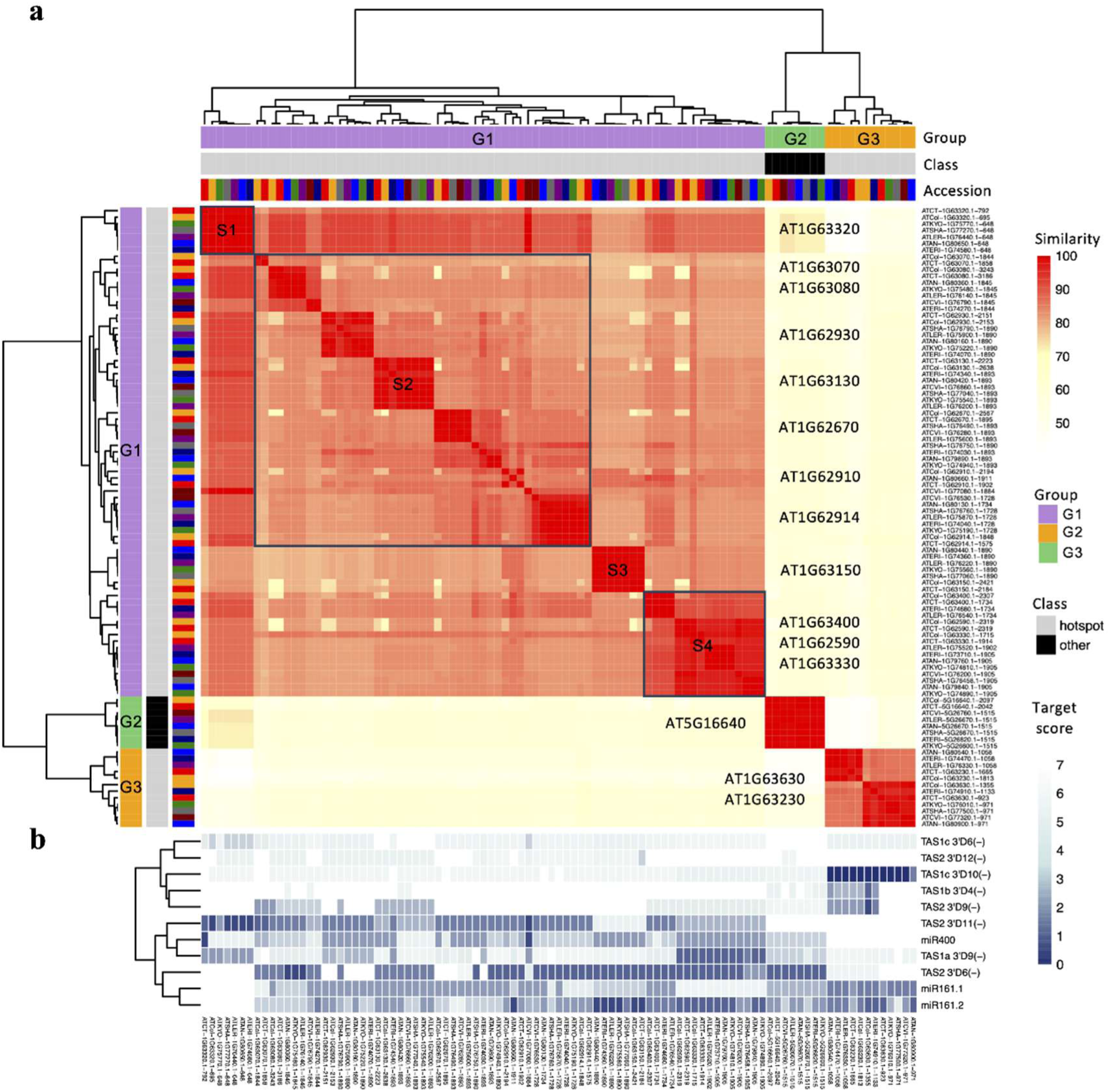
Potential gene duplication in the siRNA-producing *PPR* gene cluster of *A. thaliana*. **a.** Nucleotide sequence similarity of genes within the siRNA-producing *PPR* cluster on Chromosome 1. Eighty-seven siRNA-producing *PPR* genes located in the “hotspot” gene cluster were examined for sequence similarity across the eight *A. thaliana* accessions. Eight homologs of the siRNA-producing *PPR AT5G16640* (Col-0), which is located on Chromosome 5 but phylogenetically clustered with the hotspot siRNA-producing *PPR*s, were included for comparison. These genes form three groups: G1 (n=75), G2 (n=8), and G3 (n=12). The G1 and G3 genes are located in the “hotspot” cluster. G1 can be further grouped into four sub-groups, named S1-S4. **b.** Target site prediction of 11 sRNA triggers in the *PPR*s. The heat map colors represent targeting scores of each sRNA on each gene.

Seventy five genes in G1 share significant sequence similarity. These genes can be further classified into four sub-groups (S1-S4). Notably, homologs of the Col-0 gene *AT1G63320* form the sub-group S1, which shares a sequence similarity of > 78% with all the other genes within G1 (**Fig. 2a**). Furthermore, *AT1G63320* contains a single intron (77 nt), which is present within the coding sequences of most other siRNA-producing *PPR*s (**Fig. S2a**). These intriguing observations indicate that *AT1G63320* may represent an ancestor-like sequence that has contributed to the expansion and diversification of siRNA-producing genes within this gene cluster through duplication events.

We further investigated how the dynamics within the siRNA-producing *PPR* cluster may influence siRNA production. For this purpose, homologs of the 11 sRNAs previously shown to trigger *PPR*-siRNA production in Col-0 ^20,32^ were first identified in the other *A. thaliana* accessions and then used to predict target sites in *PPR*s in each accession. These sRNAs are highly conserved, with nearly 100% sequence identity, in all the eight accessions (**Fig. S2b; Table S4**). This analysis revealed distinctive patterns in siRNA-producing *PPR*s that belong to different groups with genes within one group sharing a similar sRNA targeting pattern (**Fig. 2b**). Interestingly, the *AT1G63320* homologs contain a highly conserved target site of TAS2 3′ D11(-), which is also present in all the G1 gene members but absent in members of G2 and G3. On the other hand, target sites of miR161.1, miR161.2 and TAS2 3′ D6(-) are absent in the *AT1G63320* homologs but prevalently adopted in the other siRNA-producing *PPR*s. This is consistent with the notion that *AT1G63320* may resemble an ancestral sequence from which the G1 group of siRNA-producing *PPR*s were derived to form a gene cluster. These additional target sRNA sites, especially miR161 and TAS2 3′ D6(-) may be responsible for the highly efficient production of siRNAs in these loci (**Fig. 2b**).

### Sequence diversification in siRNA-producing *PPR*s in the hotspot cluster

The observations indicating a higher dynamic in the siRNA-producing *PPR* cluster prompted us to quantify nucleotide sequence diversity in these genes. To determine sequence diversity, we calculated values of Pi, Watterson’s theta, and Distance (genetic distance), which led to the same conclusions (**Fig. 3a-3b**; **Fig. S3a-3b**). We compared sequence diversity among hotspot siRNA-producing *PPR*s with non-hotspot-siRNA-producing *PPR*s and non-siRNA-producing *PPR*s. To validate our analyses, the immune receptor *NLR* genes, which are the most diverse gene family in *A. thaliana* ^30,33^, were also examined. A small subset of *NLR* genes can also spawn siRNAs. Therefore, siRNA-producing and non-siRNA-producing *NLR*s were analyzed as separate groups to evaluate whether siRNA production may influence *NLR* diversification. In addition, a random set of 500 genes was used as a control group.

**Figure 3.**
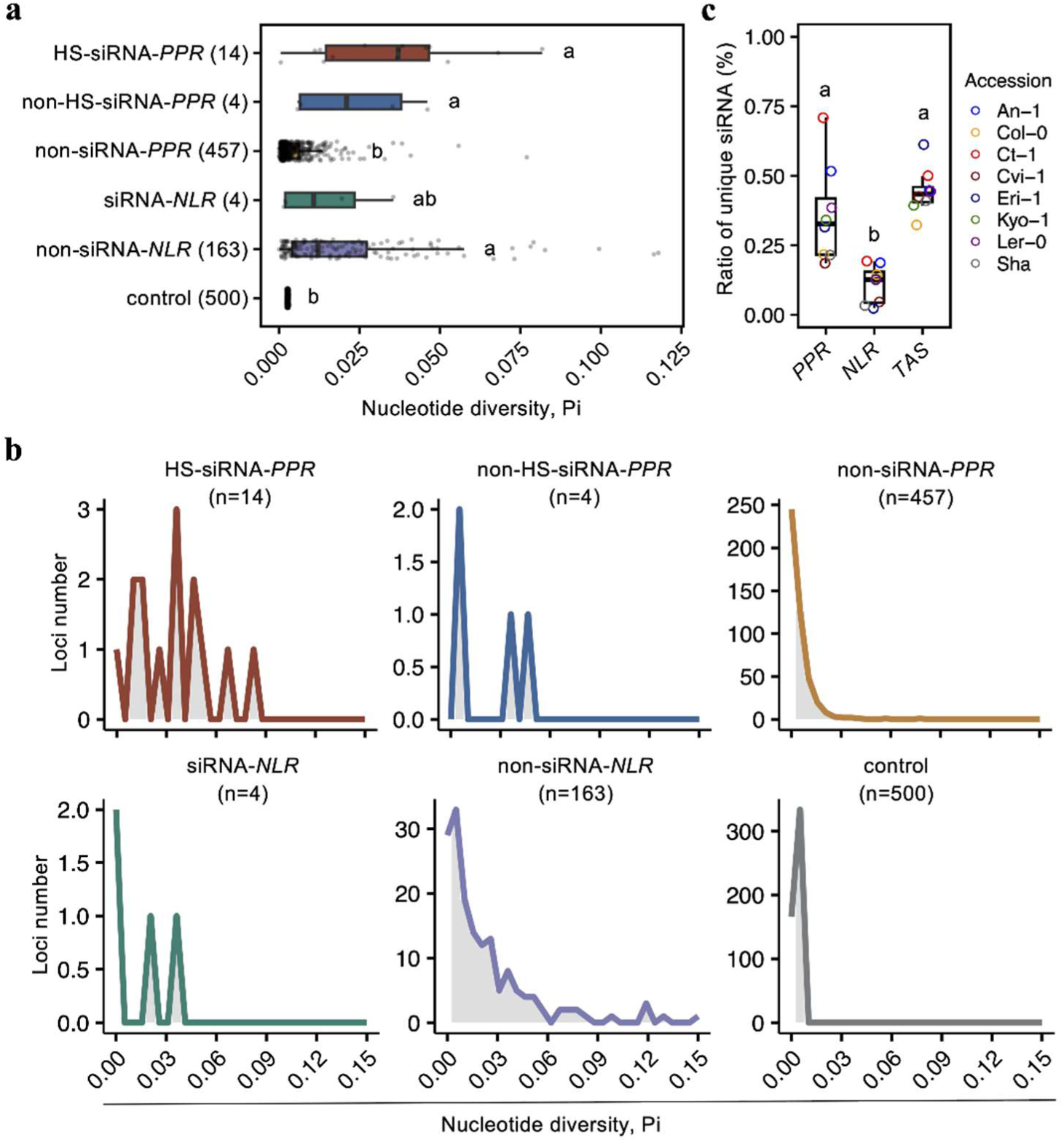
siRNA-producing *PPRs* exhibit accelerated sequence diversification in *A. thaliana*. **a.** Nucleotide diversity (Pi) was determined using sequences in eight *A. thaliana* accessions for six gene groups: hotspot-siRNA-*PPR*s (n=14), non-hotspot-siRNA-*PPR*s (n=4), non-siRNA-*PPR*s (n=457), siRNA-*NLRs* (n=4), non-siRNA-*NLR*s (n=163), and control (n=500). “n” represents the number of genes in Col-0. The ‘Kruskal-Wallis test’ method was used to compare the distribution of nucleotide diversity (Pi) among different gene groups. The ‘Bonferroni’ method was applied for multiple-comparison correction. Different letters label statistically significant differences. HS = hotspot. **b.** Distribution of Pi values of the six gene groups. **c.** Diversity of siRNAs derived from *PPR*, *NLR* or *TAS* transcripts. Distribution of siRNA diversity was calculated by ratio of unique 21-nt siRNAs produced from each gene group in the total unique siRNA population in eight *A. thaliana* accessions. Different letters label statistically significant differences.

We found the control group and the non-siRNA-producing *PPR*s exhibited the lowest genetic diversity, with median Pi values of 0.0027 and 0.0023, respectively (**Fig. 3a**). The distribution of the Pi values also displayed a similar pattern for these two gene groups, indicating a high sequence conservation (**Fig. 3b**). In contrast, both hotspot siRNA-producing *PPR*s and non-hotspot-siRNA-producing *PPRs* showed a significantly higher diversity comparable to non-siRNA-producing *PPR*s, with the median Pi values of 0.037 and 0.021. The Pi values also have wider distributions, suggesting an elevated sequence diversity in the siRNA-producing *PPR*s regardless their genomic location (**Fig. 3a-3b**). Consistent with previous studies, *NLR* genes exhibited a significantly higher diversity than the control group, with a median Pi value similar to the siRNA-producing *PPR* genes. However, no difference was observed between the siRNA-producing and non-siRNA-producing *NLR* gene groups (**Fig. 3a-3b**), indicating that, unlike *PPR*s, sequence diversification of *NLR*s is not driven by siRNA production. These observations are aligned with the different functions of *PPR*-siRNAs and *NLR*-siRNAs in host-pathogen interaction: while *PPR*-siRNAs target pathogen genes, *NLR*-siRNAs fine-tune the expression of endogenous *NLR* gene expression. Therefore, only siRNA-producing *PPR*s showed the trend of rapid evolution under selective pressure.

Despite the diversity of siRNA-producing *PPR*s, target sites of sRNA triggers have been retained to ensure siRNA production (**Fig. S3c; Table S4**), indicating that an accelerated diversification of siRNA-producing *PPR*s may lead to diversity in *PPR*-siRNAs. To test this, we compared the sequence diversity of siRNAs spawned from *PPR* loci with those derived from *NLR* or *TAS* loci. The median values of the diversity of *PPR*-siRNAs and *TAS*-siRNAs were significantly higher than that of *NLR*-siRNAs (**Fig. 3c**). Indeed, the overall diversity of 21-nt siRNA population in each *A. thaliana* accession is largely attributed to siRNAs derived from *PPR* and *TAS* loci (**Fig. S3d**). This is consistent with a role of *PPR*- and *TAS*-siRNAs in silencing gene targets from invading pathogens, thus engaging in a host-pathogen arms race ^2^.

### Contribution of high-impact variations to the diversity of siRNA-producing *PPR*s

High-impact variations include potentially functional variants, such as sequence changes that disrupt the translational reading frame (frameshift) and/or result in a premature stop codon (stop gained) ^34^. To further explore the impact of genetic variations on the diversity of siRNA-producing *PPR*s, we extended our analysis to include 1,135 *A. thaliana* genomes and detected high-impact variations from *PPR* genes. Again, *NLR* genes were included as a comparison and a random set of 500 genes was used as the control. siRNA-producing and non-siRNA-producing *PPR* and *NLR* genes were analyzed as separate groups to provide insights into a potential link between siRNA production and the presence of high-impact variation. Density of high-impact variants was determined by calculating the proportion of sequence variants with frameshift or stop gained for each gene in each group. Our results show that the control group was absent from high-impact variants, and the non-siRNA-*PPR*s were also highly conserved with a median value of 0 (**Fig. 4a**). Compared with non-siRNA producing-*PPRs*, the density of high-impact variants was significantly higher in the hotspot siRNA-producing *PPR*s (**Fig. 4a**). Non-hotpot-siRNA-producing *PPRs* also had a higher median value of 0.00222, although not statistically different from non-siRNA *PPR*s (**Fig. 4a**). Both siRNA-producing and non-siRNA-producing *NLR* genes showed the highest density of high-impact variants. These results indicate a potential contribution of gene clustering to promoting diversification of siRNA-producing *PPR*s.

**Figure 4.**
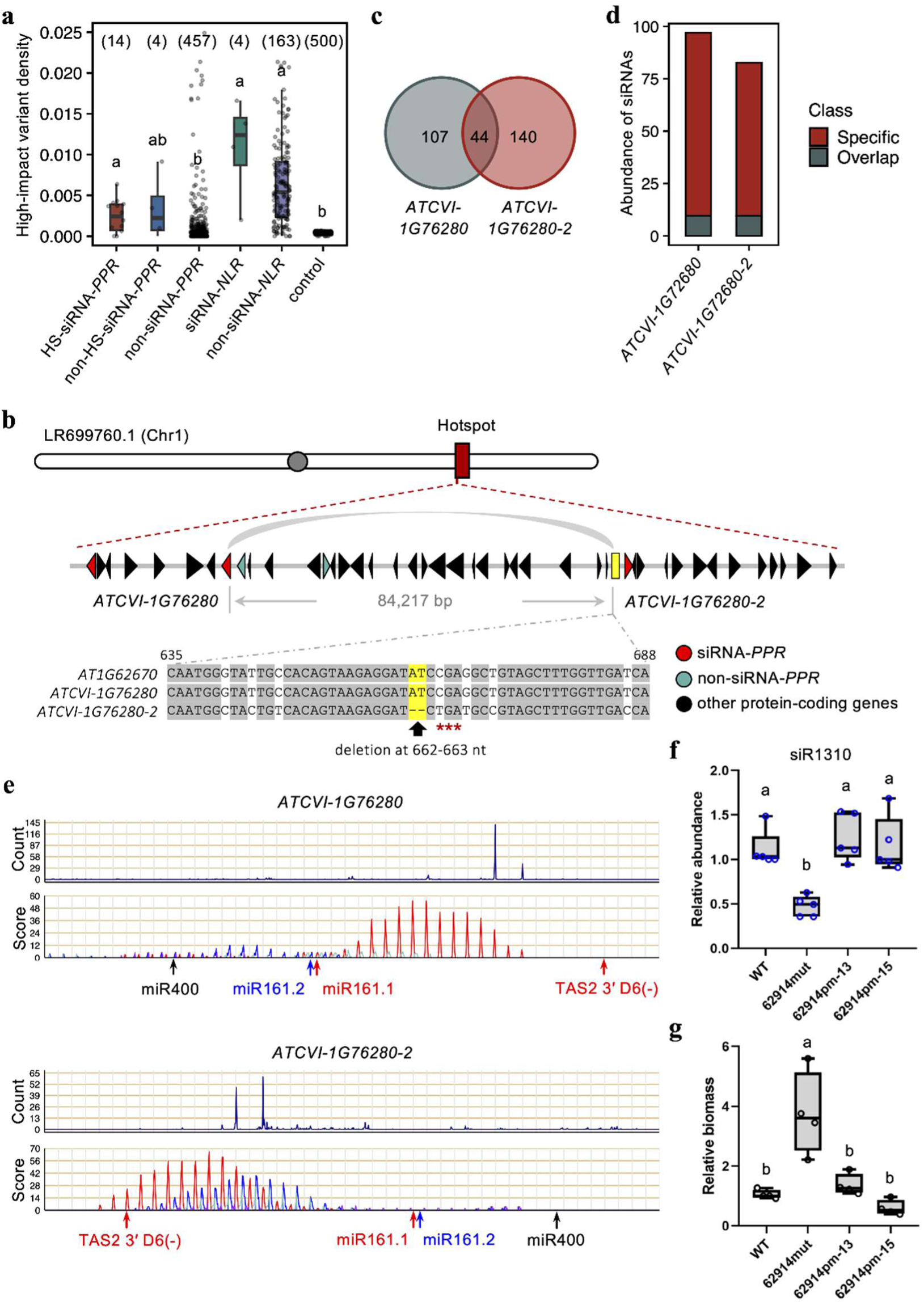
siRNA-producing *PPRs* contain a high density of high-impact variations. **a.** High-impact variation density was determined in hotspot-siRNA-*PPR*s, non-hotspot-siRNA-*PPR*s, non-siRNA-*PPR*s, siRNA-*NLRs*, non-siRNA-*NLR*s, and control groups in the pangenome of *A. thaliana*. “n” represents the number of genes in each group in Col-0. The density of high-impact variants was determined as the proportion of sequence variants with frameshift or stop gained within coding sequence for each gene in each group. Different letters label statistically significant differences. **b.** Pseudogenization of a hotspot-siRNA-producing *PPR* in Cvi-1. A region within the *PPR*-siRNA hotspot cluster in Chromosome 1 included two homologs of the Col-0 *PPR AT1G62670*. Red, blue, and black triangles represent siRNA-producing *PPR*, non-siRNA-producing *PPR*, or non-*PPR* protein-coding genes, respectively. One homolog, *ATCVI-1G76280*, is predicted to encode a PPR protein, while the other, *ATCVI-1G76280-2*, is a pseudogene located in an intergenic region (presented as a yellow rectangle). Sequence alignment shows a deletion of two nucleotides in *ATCVI-1G76280-2* results in a premature stop codon. **c.** A limited number of 21-nt siRNAs produced from both *ATCVI-1G76280* and *ATCVI-1G76280-2*. **d.** Normalized abundance (TPM) of siRNAs that are shared or specifically produced from *ATCVI-1G76280* and *ATCVI-1G76280-2*. **e.** Phasing patterns of siRNAs produced from *ATCVI-1G76280* and *ATCVI-1G76280-2*. Arrows indicate the cleavage sites of sRNA triggers, which are conserved in the two homologs. **f.** Relative abundance of siR1310 in *A. thaliana* plants determined by stem-loop PCR. WT: Col-0; *62914mut*: a T-DNA mutant of *AT1G62914*, which is major producer of siR1310; *62914pm-13* and *62914mut-15*: two independent complementation lines of *62914mut* using a CDS sequence of *AT1G62914* that contains a premature stop codon (by introducing a T-to-A point mutation at the nucleotide position 432). Values are mean ± s.d. of three biological replicates. One-way ANOVA with post hoc Tukey testing were used for statistical analysis. Different letters label statistically significant differences (*p* < 0.05). **g.** Relative biomass of *Phytophthora capsici* in inoculated plants evaluated using qPCR at 3 days post inoculation (dpi). Values are mean ± s.d. of three biological replicates. Different letters label significant differences (*p* < 0.05).

From the analysis of high-impact variations, we detected instances of pseudogenization in hotspot siRNA-producing *PPR* genes. Although these transcripts are inactivated for protein-coding ability, they continue to spawn siRNAs. One such instance involves homologs of the Col-0 gene *AT1G62670*, which has two homologous sequences in the accession Cvi-1. Both sequences are located within the *PPR*-siRNA hotspot region in Chromosome 1 of Cvi-1 and share similarities of 97.79% and 90.50% with Col-0 *AT1G62670*, respectively (**Fig. 4b**). One of the homologs, *ATCVI-1G76280*, is predicted to encode a PPR protein, while the other, named *ATCVI-1G76280-2*, represents a non-coding locus in an intergenic region of the plus strand. These two loci are situated 84,217 bp apart. *ATCVI-1G76280-2* appears to have experienced one AT deletion at the nucleotide position 662-663 of *ATCVI-1G76280*, resulting in premature stop codon (**Fig. 4b**). Importantly, both Cvi-1 loci spawn siRNAs; however, due to sequence diversification, they produce distinct populations of siRNAs with specific sequences (**Fig. 4c**). Less than 10% of the siRNA populations was shared by both loci (**Fig. 4d**). Target prediction of these two loci revealed potential target sites of the same sRNA triggers. Importantly, abundant siRNAs were produced from similar regions between the target sites of TAS2 3′ D6 (-) and miR161.1 in both loci (**Fig. 4e**). Phasing patterns of the siRNAs are consistent with TAS2 3′ D6 (-) and miR161.1 being the major triggers for siRNA production in both loci, indicating a preservation of siRNA production mechanism despite gene duplication and pseudogenization.

A similar example was found in the accession Sha. In this case, two loci share sequence similarity with the Col-0 gene *AT1G63080* (**Fig. S4**). One of the homologous sequences has a premature stop codon due to a C-to-G point mutation at the nucleotide position 922. Both loci produce siRNAs, but their siRNA populations are highly specific due to sequence divergence. Nevertheless, siRNA productions from these two loci are triggered by the same sRNAs with similar phasing patterns (**Fig. S4**). Taken together, these observations suggest that an accelerated diversification of *PPR*-siRNA-producing sequences could be achieved through sequence diversification, gene duplication, and high-impact variation including pseudogenization.

Given that *PPR* transcripts could potentially produce both proteins and siRNAs, we conducted experiments to clarify the role of siRNA production in the defense contribution of *PPR* genes. *PPR*-siRNAs have been previously reported to enhance *A. thaliana* defense against the oomycete pathogen *P. capsici*, likely by silencing target gene(s) in the invading pathogen ^14^. In particular, siR1310 and siR1310-like *PPR*-siRNAs (collectively called siR1310 hereafter) are predicted to silence the *P. capsici* gene *Phyca_554980* and artificial introduction of siR1310 into *P. capsici* significantly reduced the virulence in *A. thaliana* ^14^. Furthermore, an *A. thaliana* mutant of a siR1310-producing *PPR*, *AT1G62914*, exhibited increased susceptibility to *P. capsici* ^14^. We generated complementation lines in the *at1g62914* mutant background by introducing a mutated sequence of *AT1G62914* (*62914pm*), which contains a T-to-A point mutation at the nucleotide position 432 to implement a premature stop codon (**Fig. S5a; Table S5**). Although the full-length mRNAs could still be produced in the complementation lines, western blotting failed to detect protein products, likely due to the premature stop codon (**Fig. S5b-c**). We confirmed that the production of siR1310 was restored in the *62914pm* complementation lines to the wildtype level (**Fig. 4f**). Furthermore, the *62914pm* construct also restored the hypersusceptibility phenotype of the *at1g62914* mutant to *P. capsici* (**Fig. 4g**). These results suggest that pseudogenized *PPR* loci could still contribute to plant defense, highlighting the functional significance of siRNA production of *PPR* genes in plant defense.

### siRNA-producing PPRs form an ancient clade that exhibits within-species diversification

PPR proteins are found in all eukaryotes with an expansion in plants ^16^. Previously, siRNA production from *PPR* transcripts has been observed in 32 dicot plants ^18,35,36^. We examined *PPR*-siRNA production in 55 plant species, including three early plants, ten monocots, and 42 eudicots (**Fig. S6a**). These species were chosen based on the availability of high-quality reference genome, gene annotations, and sRNA-seq dataset (**Table S5**). To accommodate variation in genome annotations and updates in genome assemblies, PPR proteins from each genome were re-annotated based on sequence similarity using blastp against previously identified PPR proteins (https://ppr.plantenergy.uwa.edu.au/) ^25^. In general, each species has at least one gene encoding each of the four PPR subfamily proteins, and up to 150 DYW class, 489 E class, 589 P class and 322 PLS class proteins (**Table S6**).

To detect common themes in siRNA-producing *PPR*s in these evolutionarily diverse species, 21-nt siRNAs were mapped to a specific *PPR* locus in each species. A *PPR* locus was considered to produce siRNAs using a cutoff of normalized TPM > 5. *PPR*-siRNAs were detected from all but five species, including two dicots, two monocots and one early plant. In most species, the majority of PPR-siRNAs are spawned from P class PPR-encoding genes (**Fig. S6b; Table S6**). These results indicate that *PPR*-siRNA production from P class PPR-encoding genes is an evolutionarily conserved phenomenon in plants.

We further investigated the evolutionary relationship of the siRNA-producing PPRs. A phylogenetic tree of 11,617 PPR sequences from 17 representative species spanning major land plant lineages, including monocots and eudicots, was generated (**Fig. 5a-5b**; **Table S7**). We incorporated 168 PPRs that are evident to produce siRNAs from publicly available databases into tree, which revealed that 122 of the siRNA-producing PPRs are clustered in a single clade (**Fig. 5b**; **Table S8**). This remarkable result suggests that siRNA production is an ancient function of this specific PPR subfamily. We further examined the 546 PPRs from this “siRNA-producing” clade, color-coded by the species of origin. This analysis revealed a strong pattern of within-species diversification (**Fig. 5c**) in this clade, which is consistent with their role in plant defense. Interestingly, we found a monocot clade, with 22 PPRs from rice and maize, paraphyletic to the dicot clade containing siRNA-producing PPRs. Although these monocot PPRs are not known to produce siRNAs in the currently available dataset, it remains to be confirmed whether they retain this activity.

**Figure 5.**
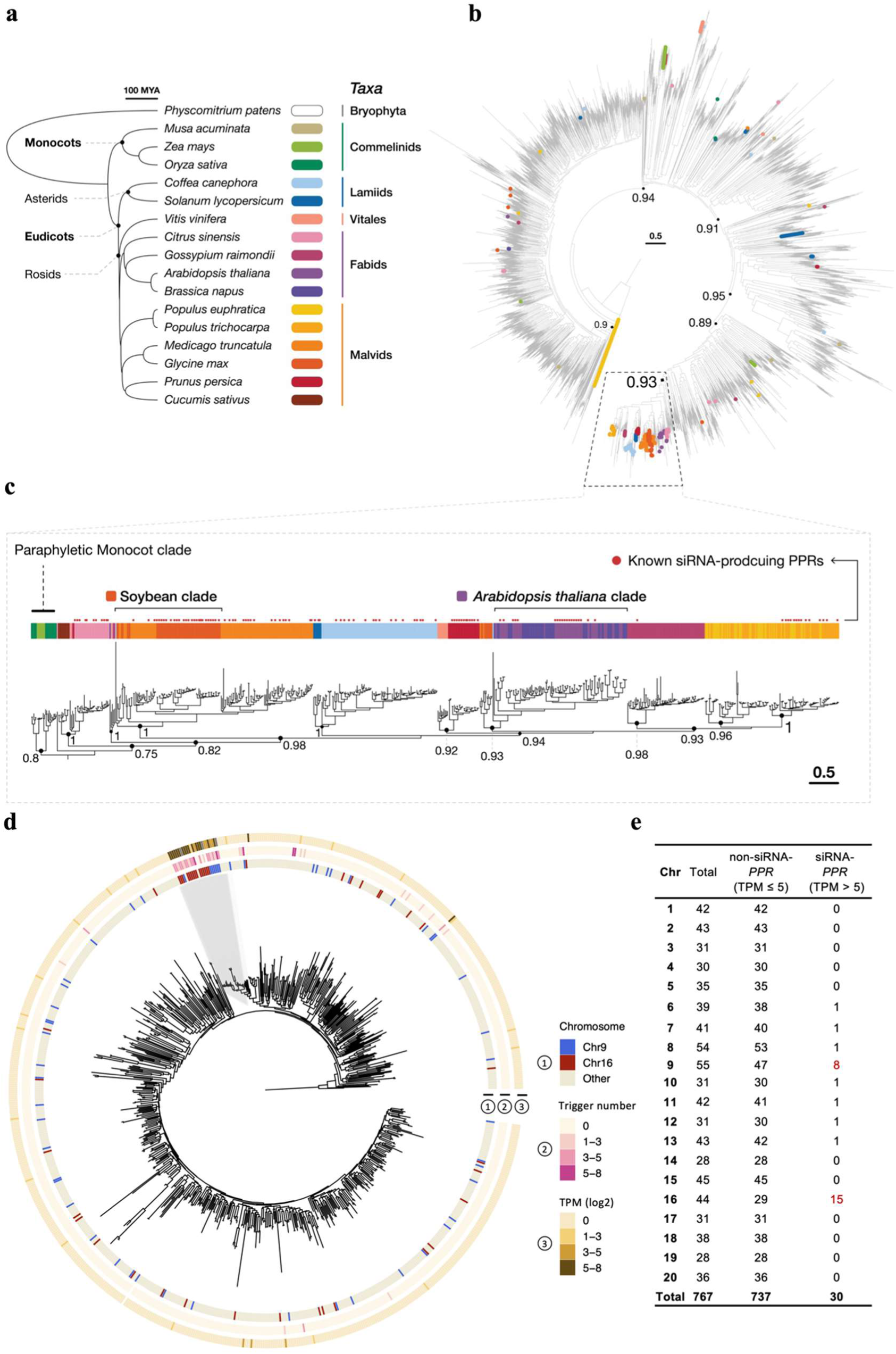
siRNA-producing PPRs form an evolutionarily conserved monophyletic clade. **a**. Taxonomic tree of 17 representative species across land plants. **b**. Phylogenetic tree of 11,617 PPR sequences from 17 plant species. 168 sRNA-producing PPRs are highlighted by red dots. Four PPRs, including CCM1_CANAW from *Candida albicans*, CCM1_SCHPO from *Schizosaccharomyces pombe*, and LPPRC from both *Homo sapiens* and *Rattus norvegicus*, are used as the outgroups. **c**. Phylogenetic tree of 546 PPRs belonging to the siRNA-producing clade with the 122 known siRNA-producing PPRs highlighted by red dots. The PPRs are colored by the species of origin. Numbers next to the nodes shown on the phylogenetic trees indicate bootstrap value. **d.** Phylogenetic tree of the entire complement of 767 soybean PPRs based on amino acid sequences. Ring 1 indicates the chromosomal location of the gene. Ring 2 indicates the number of miRNA trigger target sites predicted from each *PPR*. Ring 3 presents the abundance of 21-nt siRNAs, represented by log_2_-tranformed TPM, derived from each *PPR* in soybean leaves. **e.** Number of non-siRNA-producing and siRNA-producing *PPR* genes in each chromosome of soybean.

We next examined whether the siRNA-producing *PPRs* are also physically clustered in other species as observed in *A. thaliana*. For this purpose, soybean (Genome assembly, Glycine_max_v2.1), which has 767 PPR genes, was analyzed (**Table S9**). In total, we identified 30 *PPRs* that produce 21-nt siRNAs. Twenty-four of the siRNA-producing genes putatively encode P class PPR proteins and belong to a single phylogenetic clade (**Fig. 5d**). These genes are responsible for the production of 84.85% of the total 21-nt *PPR-*siRNA population in soybean (**Fig. 5d**; Ring 3). Long branches in the phylogenetic tree of this clade indicates intra-species diversification of the genes. Furthermore, members in this phylogenetic clade are prominent targets of sRNAs that can trigger *PPR*-siRNA production (**Fig. 5d**; Ring 2). All these features are reminiscent to what we have observed from the siRNA-producing *PPR*s in *A. thaliana*.

Interestingly, not only the major soybean siRNA-producing *PPR*s form a single phylogenetic group, they are also physically linked as two clusters (**Fig. 5d**; Ring 1; **Fig. 5e**; **Table S9**): seven of the eight siRNA-producing *PPR*s on Chromosome 9 are located within a ∼8 Mb region (nucleotides 39,625,286 – 47,564,658) and the 15 siRNA-producing *PPR*s on Chromosome 16 are located within a ∼ 6 Mb region (nucleotides 29,727,924 – 36,087,973). Soybean is a tetraploid through whole genome duplications ^37^. The two siRNA-producing *PPR* gene clusters may be generated from the duplication event. These results indicate that, like in *A. thaliana*, soybean also employs designated *PPR* gene clusters as hotspots for siRNA production.

### Heterologous expression of *PPR*-siRNAs in *N. benthamiana* enhances disease resistance

Considering *PPR*-siRNA production as an evolutionarily conserved process in plants, we explored how plant defense can be elevated by enhancing production of these antimicrobial agents. As a proof-of-concept, we designed a *PPR*-siRNA-producing cassette by co-expressing sRNA trigger(s) and a *PPR*-siRNA-producing sequence from *A. thaliana* in *N. benthamiana*. To exclude potential involvement of protein product(s), *AT1G62914pm* was used as the siRNA producing sequence. Target prediction in *AT1G62914pm* using 158 known miRNAs produced by *N. benthamiana* ^38^ did not retrieve any significant hits. Therefore, we co-expressed *AT1G62914pm* with sRNA triggers from *A. thaliana*, in particular miR161 and TAS2 3′ D6 (-), which are associated with the production of some of the highest siRNA-producing *PPR*s. As a control, we generated a variant of *AT1G62914pm*, referred to as *AT1G62914del* (**Fig. S7**; **Table S5**), which contains a deletion of a 159 bp (631 to 789 nt) fragment from which most siRNAs, including siR1310, are produced in wildtype *AT1G62914*. Importantly this mutant gene still retains the target sites of the sRNA triggers.

When *AT1G62914pm* was co-expressed with both miR161 and TAS2 3′ D6 (-), but not miR161 alone, we detected an increased accumulation of *PPR*-siRNAs in *N. benthamiana*, as indicated by an elevated level of siR1310 (**Fig. 6a**). In contrast, co-expression of the trigger sRNAs with *AT1G62914del* did not increase siR1310 accumulation. We then inoculated the *N. benthamiana* leaves using zoospore suspension of *P. capsici*. siR1310 was previously shown to silence the *P. capsici* gene *Phyca_554980*. Consistent with the increased accumulation of siR1310 in tissue co-expressing *AT1G62914pm*, miR161 and TAS2 3′ D6 (-), we observed reduced transcript levels of *Phyca_554980* (**Fig. 6b**) and enhanced resistance to *P. capsici*, reflected by reduced lesion sizes (**Fig. 6c-6d**) and pathogen biomass (**Fig. 6e**). These results demonstrate that engineering *PPR*-siRNAs can be employed as a promising immune enhancement strategy.

**Figure. 6.**
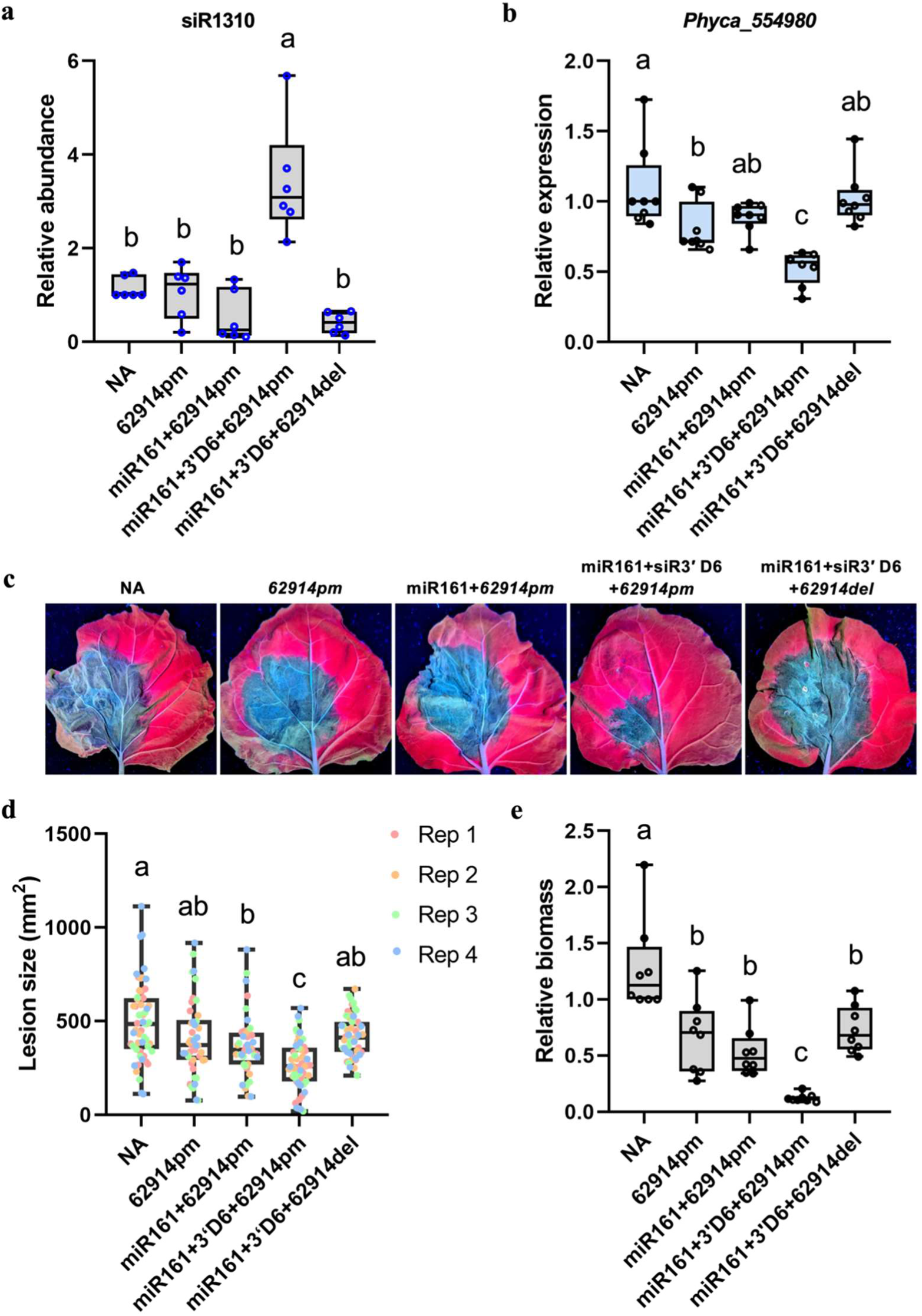
Ectopic expression of a *PPR*-siRNA-producing cassette in *N. benthamiana* enabled siRNA accumulation and enhanced resistance to *P. capsici*. a. Relative abundance of siR1310 in *N. benthamiana* leaves determined by stem loop qRT-PCR. Different combinations of *Agrobacterium* carrying the *A. thaliana PPR* gene *AT1G62914pm* (carrying a premature stop codon) with sRNA triggers miR161 and/or TAS2 3’D6(-) were used to infiltrate *N. benthamiana* leaves. *P. capsici* zoospores were inoculated at 24 hours after Agro-infiltration and leaf tissues were collected at 3 days post *P. capsici* infection (dpi) for further analyses. NA: no Agro-infiltration; *62914del*: *AT1G62914pm* with a 159 bp deletion (631-789 nt) that eliminate siR1310 production without affecting the target sites of sRNA triggers. Values are mean ± s.d. of three biological replicates. One-way ANOVA with post hoc Tukey testing was used for statistical analysis. Different letters label statistically significant differences (*p* < 0.05). **b.** Transcript abundances of the *P. capsici* gene *Phyca_554980* determined by qRT-PCR. Values are mean ± s.d. of five biological replicates. One-way ANOVA with post hoc Tukey testing was used for statistical analysis. Different letters label statistically significant differences (*p* < 0.05). **c.** Disease symptoms of *N. benthamiana* leaves at 3 dpi. **d.** Lesion sizes in the inoculated *N. benthamiana* leaves measured at 3 dpi. At least seven leaves per treatment were examined across three replicates. One-way ANOVA with post hoc Tukey testing was used for statistical analysis. Different letters label statistically significant differences (*p* < 0.05). **e.** Biomass of *P. capsici* in inoculated *N. benthamiana* leaves determined at 3 dpi. Values are mean ± s.d. of five biological replicates. One-way ANOVA with post hoc Tukey testing was used for statistical analysis. Different letters label statistically significant differences (*p* < 0.05).

## Discussion

The secondary siRNA pathway has emerged as a major contributor to plant defense. This defense function could be attributed to two different mechanisms: (i) siRNAs produced from *NLR* and *TAS3* transcripts might regulate the expression of their endogenous targets to fine-tune immune-related gene expression ^8–13^; (ii) siRNAs produced from *PPR* and *TAS1/TAS2* transcripts may silence target genes in invading eukaryotic pathogens through host-induced gene silencing or HIGS ^14^. Unlike endogenous gene targets, pathogen transcripts targeted by plant sRNAs are presumed to be under selective pressure to evade HIGS ^19^. Therefore, siRNA-source sequence that directly target pathogen genes may have different evolutionary trajectories from those regulating endogenous targets.

In this study, we investigated the evolutionary features of PPR genes that give rise to siRNAs. PPRs represent one of the largest protein families in land plants ^16^. As proteins, they are known to regulate RNA metabolism in mitochondria and chloroplasts ^17^. We divided PPRs into two classes based on whether they produce siRNAs and demonstrate that siRNA-source *PPR*s exhibit a remarkable sequence diversity, which is in stark contrast to the highly conserved non-siRNA-source *PPR*s. The siRNA-producing PPRs also belong to the “Restorer of Fertility-Like” (RFL) group, although not all RFLs are siRNA-producers. In *A. thaliana* Col-0, 13 out of 26 RFLs produce siRNAs ^39^. RFLs were previously reported to exhibit intraspecific diversity, which was attributed to their role in regulating organelle gene expression ^39,40^. The strong correlation between sequence diversification and the ability to produce siRNAs suggests that host-pathogen co-evolution is an important factor that specifically impacts the evolution of siRNA-producing PPRs.

Through a survey of 55 plant species encompassing dicots, monocots and early plants, we found siRNAs derived from *PPR* sequences in most species, expanding the previous study suggesting that *PPR*-siRNA production is a conserved phenomenon in eudicots^18^. Remarkably, the siRNA-producing PPRs form a monophyletic group, strongly suggesting that siRNA production is their ancient function. Furthermore, genes within this group exhibit a high degree of within-species diversification, consistent with a role as a new class of evolutionarily conserved defense genes.

In both *A. thaliana* and soybean, the majority of siRNA-producing PPRs are physically clustered, presumably promoting sequence diversification. Genes encoding the NLR immune receptors often form multi-gene clusters, which are believed to contribute to the rapid expansion and diversification of pathogen recognition capacity ^41–43^. Similarly, the physical clustering may contribute to generating diversity in the siRNA-producing *PPR*s. In particular, we found that the majority of the siRNA-producing *PPR*s located within the cluster on Chromosome 1 in *A. thaliana* are potentially derived from a single gene (*AT1G63320* in Col-0), which shares a high sequence similarity (>78%) to all other siRNA-producing *PPRs*. It was proposed that a few *PPR* genes with introns in *A. thaliana* and rice are probably ancient and served as templates for the numerous intron-less genes created during the retrotransposition-driven expansion of the *PPR* gene family ^44^. Notably, *AT1G63320* is the shortest gene (∼ 700 nucleotides) in this cluster. Furthermore, it contains a single intron (77 nt) that is present within the coding sequences of most other *PPR*s within this cluster. Therefore, an *AT1G63320*-like sequence could serve as an ancestor that leads to the expansion of this siRNA-producing gene cluster. Members in this cluster may then undergo further sequence diversification.

*PPR* genes can function through their encoded protein products or derived siRNAs. We found multiple examples of pseudogenization events within the siRNA-producing *PPR* cluster in different *A. thaliana* accessions that, although not encoding functional proteins, preserve the trigger sRNA target sites and maintain the ability to produce siRNAs. Functional analysis confirmed that a pseudogenized *PPR* can still confer defense, showing that their main biological function is through producing anti-microbial siRNAs. Often, the pseudogenized gene produces a largely diverged siRNA population from its functional homolog, indicating high-impact variations such as pseudogenization contribute to the generation of a wide variety of siRNAs, which is beneficial for the plants to target multiple pathogen genes ^19,45^.

In *A. thaliana*, secondary siRNAs have been implicated in contributing to defense to the oomycete pathogen *P. capsici*, and the fungal pathogens *Botrytis cinerea* and *Verticillum dahliae* ^14,46^. Interestingly, both oomycete and fungal pathogens produce effector proteins that can suppress the secondary siRNA accumulation ^14,47^, suggesting a host-pathogen arms race centered on this defense-related pathway. In particular, the *Phytophthora* effector PSR2 suppresses the biogenesis of secondary siRNAs derived from *TAS1*, *TAS2*, and *PPR* transcripts in *A. thaliana* ^2^. Therefore, secondary siRNAs contribute to broad-spectrum defense against a variety of microbial pathogens.

Our finding that the siRNA-producing PPRs belong to an evolutionarily conserved clade supports the possibility of engineering siRNA-producing sequences in a wide range of plants to enhance disease resistance. We conducted a proof-of-concept study by introducing a *PPR*-siRNA-producing cassette using corresponding sequences in *A. thaliana* into *N. benthamiana*. Expressing this cassette led to a significant accumulation of specific *PPR*-siRNAs and elevated resistance to *P. capsici*. Importantly, siRNA production required co-expression of the siRNA-producing sequence with two sRNA triggers. In contrast, the well-established trigger sRNA miR161, by itself, is insufficient to initiate *PPR*-siRNA production. This is consistent with the observation that almost all the siRNA-producing *PPR*s possess predicted target sites of multiple different sRNA triggers. Additional experiments are required to further optimize the siRNA production efficiency by exploring different combinations of trigger sRNAs with *PPR* genes. It will also be exciting to test the system in diverse crops.

In summary, this study identified a specific *PPR* cluster that has evolved to produce siRNAs conferring disease resistance. The unique evolutionary features of this siRNA-producing gene clade highlight accelerated diversification, potentially driven by co-evolution with pathogens. Our analysis of a specific *PPR* family involved in defense opens new opportunities to develop broad-spectrum disease resistance in a wide range of crops. Bespoke siRNAs that efficiently target selected pathogen gene(s) may be achieved by engineering the siRNA-producing *PPR* sequences.

## Methods

### Reference genome and gene annotation sources

The genome sequence (version TAIR10) and gene annotation (version Araport11) of *A. thaliana* Col-0 accession were obtained from EnsemblPlant v56 ^48^. The Ct-1 genome was sequenced using Illumina paired-end reads, and both its genome sequence and gene annotation were derived from the 19 genomes project ^22^. The genome sequences of An-1, Cvi-1, Eri-1, Kyo-1, L*er*-0, and Sha were obtained through high resolution PacBio sequencing ^23^ and the annotations were originated from the 1001 genomes project ^49^. The genome sequences and gene annotations for 54 additional plant species were retrieved from public databases, including NCBI (https://www.ncbi.nlm.nih.gov/), EnsemblPlant v56, and Phytozome v13 ^50^ (**Table S6**).

For *A. thaliana* Col-0, the annotations of eight *TAS* loci (*TAS1a/b/c*, *TAS2*, *TAS3a/b/c* and *TAS4*) and 27,655 protein-coding genes were obtained from EnsemblPlant v56 ^48^. The amino acid sequences of 475 PPRs were downloaded from the “PPR” website ^51^. The 167 NLRs were annotated using the nlrsnake software (https://github.com/TeamMacLean/nlrsnake) ^52^. Homologous genes of *TAS*, *PPRs*, and *NLRs* in *A. thaliana* Col-0 in the other seven accessions were identified by the method described in the “Homologous gene identification” section.

### sRNA-seq data analysis

Rosette leaves from 3-week-old *A. thaliana* were used for sRNA libraries. The library construction was initiated from total RNAs and followed the instruction of NEBNext multiplex small RNA library prep set for Illumina (E7300S). The sRNA libraries were sequenced using the Illumina Hiseq 4000 platform with single-end reads of 150 bases. Before analysis, the raw sequencing reads in fastq format were removed the adapter sequence AGATCGGAAGAG using the software Cutadapt v3.2 ^53^. Any reads not successfully trimmed or containing more than one ‘N’ base were discarded by the adapter removal process. Quality filtering was performed by Trimmomatic v0.36 ^54^ using single-end mode with LEADING:3, TRAILING:3, SLIDINGWINDOW:4:15, MINLEN:18, and MAXLEN:28. After quality control, the resulting reads with length ranging from 18- to 28-nt were firstly mapped to known the sequences of structural RNAs from Rfam v14.9 ^55^. The categories of structural RNAs included transfer RNAs (tRNAs), ribosomal RNAs (rRNAs), small nuclear RNAs (snRNAs), and small nucleolar RNAs (snoRNAs). The remaining unmapped reads were secondly mapped to the *A. thaliana* reference genome using Bowtie v4.4.6 ^56^, allowing all alignments (-a) and zero mismatch (-v 0) per read. The successfully mapped reads represent clean reads. The diversity of siRNAs spawned from a locus in an *A. thaliana* accession was assessed using the ratio of unique 21-nt siRNAs produced from that gene to the total number of unique siRNAs from the accession.

sRNA-seq data from other plant species were processed and analyzed using the same pipeline as described above. For each locus, the sRNA abundance (transcripts per million, TPM) was normalized by dividing the number of sRNAs produced from that gene by the total number of genome-matched clean reads in a library.

### Identification of homologous genes

Identification of homologs of Col-0 protein-coding genes in other *A. thaliana* accessions was conducted using SimpleSynteny (v1.3.2) ^26^, blastn and blastp (blast+, v2.9.0) ^27^ using default parameters. 27,377 genes in Col-0, excluding those derived from mitochondria and chloroplasts, as well as CDS sequences shorter than 80 aa, were used as queries. Initial homologs were filtered based on query coverage > 30%, subject coverage > 30%, identity > 60%, and bit score > 50. The remaining homologs were filtered based on the scoring system, as shown in the following formula,

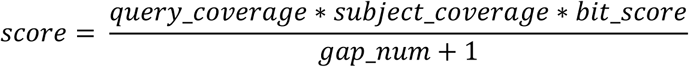

In this formula, 𝑞𝑢𝑒𝑟𝑦_𝑐𝑜𝑣𝑒𝑟𝑎𝑔𝑒 and 𝑠𝑢𝑏𝑗𝑒𝑐𝑡_𝑐𝑜𝑣𝑒𝑟𝑎𝑔𝑒 indicates the coverage of aligned query and subject sequence, respectively. 𝑏𝑖𝑡_𝑠𝑐𝑜𝑟𝑒 reflects the overall quality of an alignment. 𝑔𝑎𝑝_𝑛𝑢𝑚 represents the number of gaps in an alignment. Higher scores indicate more reliable homologies. Retaining the highest score, 26,877, 26,831, 26,753, 26,892, 26,887, 26,874, and 26,828 homologs for the 27,377 genes in Col-0 were identified in the An-1, Ct-1, Cvi-1, Eri-1, Kyo-1, L*er*-0, and Sha, respectively. Among these genes, 467, 471, 459, 465, 464, 467, and 461 were homologous to the 475 *PPRs* in Col-0, while 140, 144, 122, 140, 129, 137, and 133 were homologous to the 167 *NLRs* in Col-0.

Homologs for the non-coding genes in Col-0, including eight *TAS* loci (*TAS1a/b/c, TAS2, TAS3a/b/c,* and *TAS4*) and 11 sRNA triggers, were identified as the best alignment with the highest similarities in other seven *A. thaliana* accessions. These 11 sRNA triggers including miR161.1, miR161.2, miR400, TAS1a 3′ D9 (-), TAS1b 3′ D4 (-), TAS1c 3′ D6 (-), TAS1c 3′ D10 (-), TAS2 3′ D6 (-), TAS2 3′ D9 (-), TAS2 3′ D11 (-), and TAS2 3′ D12 (-) were obtained from both miRbase ^57^ and a previous study ^32^.

### Determination of sRNA targeting scores on *PPRs*

The GSTAr.pl algorithm (version 1.0) ^38^ was utilized to calculate the targeting score for the homologs of 11 sRNA triggers on *PPR* transcripts in all eight *A. thaliana* accessions. Genes with an Allenscore < 4 were defined as predicted targets.

### Nucleotide diversity analysis

Nucleotide diversity was analyzed for six gene groups across eight *A. thaliana* accessions. The hotspot-siRNA-*PPR*, non-hotspot-siRNA-*PPR*, non-siRNA-*PPR*, siRNA-*NLR*, non-siRNA-*NLR* groups included 14, 4, 457, 4, and 163 genes, respectively, as well as a control group consisting of a random set of 500 genes. The control group was generated by performing 1000 random selections, each containing 500 genes. Nucleotide diversity was evaluated by determining Pi, Watterson’s theta, and Distance values. For each gene, codon-based alignment (https://github.com/lyy005/codon_alignment) was conducted on the CDS sequences of homologs in eight *A. thaliana* accessions. If one or more accessions lack a specific ortholog, those accessions were excluded from the calculation for that particular ortholog to avoid skewing in assessing the diversity. The Pi and Watterson’s theta value were estimated using the sliding.window.transform function from the R package Popgenome ^58^ with a sliding window size and a moving window size of 50. The Distance value was calculated using the dist.dna function from the R package APE ^59^.

Correlation between the sequence diversity in *PPR*, *NLR*, or *TAS* genes and the diversity of siRNAs spawned from the corresponding loci was evaluated using the cor.test method in the R package STATS^60^. The diversity of siRNAs spawned from each gene in the eight *A. thaliana* accessions was assessed using the ratios of unique 21-nt siRNAs produced from that gene to the total number of unique 21-nt siRNAs from each accession. Spearman method was used to assess the association strength and direction between two ranked variables and a *p-value* < 0.05 was considered statistically significant.

### PPR identification and Phylogenetic analyses

Phylogenetic trees for PPRs in *A. thaliana* and soybean were constructed using the amino acid sequences of 475 PPRs from Col-0 accession and 767 PPRs from soybean, respectively. The PPR sequences were aligned using MUSCLE (v3.8.31) ^61^ and automatically trimmed with trimAI (v1.2rev59) ^62^. Phylogenetic trees were constructed using iqtree2 (v2.0.4) ^63^ with estimated substitution models of VT+F+G4 in *A. thaliana* and LG+F+G4 in soybean and 1000 ultrafast bootstrap replicates. The trees were visualized using the R package ggtree (v3.6.2) ^64^.

The species phylogeny was constructed using the amino acid sequences of AT2G21710 orthologs, which encode a mitochondrial transcription termination factor. The phylogenetic relationships among the 55 plant species are consistent with a previous report ^65^. The phylogenetic tree was constructed using iqtree2 with the estimated substitution model JTTDCMut+I+G4.

We identified Pentatricopeptide Repeat (PPR) proteins by searching four PFAM HMM profiles (PF01535, PF12854, PF13041, PF13812) against 17 RefSeq proteomes using hmmsearch [--max] (**Table S7**) ^66^. A total of 19,352 sequences with hits to at least one of these profiles and an e-value of 0.05 or lower were retained. To ensure data quality, we filtered sequences based on length, excluding those shorter than 150 amino acids or longer than 1,500 amino acids. After this filtration, 19,274 sequences remained. Redundant sequences were removed using the “unique” function of the BiocGenerics R package, resulting in a final dataset of 11,617 unique PPR sequences (**Table S8**) ^67^.

For phylogenetic analysis, we incorporated 168 reference sRNA-producing PPRs and four outgroup PPRs, including CCM1_CANAW from *Candida albicans*, CCM1_SCHPO from *Schizosaccharomyces pombe*, and LPPRC from both *Homo sapiens* and *Rattus norvegicus* into the already extracted PPR sequences (**Table S8**). These sequences were aligned using FAMSA v2.2.2 with the default options ^68^. The alignment was then used to construct a phylogenetic tree using FastTree v2.1.10 with the [-lg] model ^69^. This phylogenetic tree was rooted at the LPPRC sequences. The sRNA-producing clade was identified based on a well-supported branch within the phylogeny. Among the 546 sequences from this clade, 22 are from a paraphyletic clade with two monocot plant species, and 122 are known siRNA-producing PPRs. These sequences were realigned using MAFFT v7.525 ^70^ and a new phylogenetic tree was constructed using FastTree v2.1.11 ^69,70^. This phylogenetic tree was rooted at the paraphyletic clade. All phylogenetic trees were visualized using iTOL and manually curated ^71^. The taxonomic tree for the 17 selected species was retrieved from TimeTree 5 (timetree.org) ^72^.

### High-impact variation density analysis

Whole genome sequence variation data in 1,135 *A. thaliana* accessions were downloaded from the 1001 Genome Project ^73^. High-impact variants were characterized by their potential to trigger deleterious effects on gene functions, such as introducing premature stop codons through single nucleotide polymorphisms (SNPs) or causing frameshift mutations through small insertions or deletions.

High-impact effects were assessed in six gene groups, including hotspot-siRNA-*PPR*s, non-hotspot-siRNA-*PPR*s, non-siRNA-*PPRs*, siRNA-*NLR*s, non-siRNA-*NLR*s, and random genes. The high-impact density for each gene was determined by the proportion of variants carrying frameshift and stop gained within its coding sequence.

### Plant growth and *Phytophthora* infection

Seeds for this study were ordered from the Arabidopsis Biological Resource Center (ABRC) and included CS97808 (An-1), CS76786 (Ct-1), CS97810 (Cvi-1), CS97813 (Eri-1), CS97807 (Kyo-1), CS97814 (L*er*-0), CS97811 (Sha), and CS737432 (*62914mut*). Plants were grown in a growth room with controlled environment at 22 ± 2℃ and 12h light/12h dark photoperiod.

*Phytophthora capsici* isolate LT263 was cultured and used for plant inoculation by zoospore suspensions as previously described ^14^. Biomass of *P. capsici* was determined by qPCR at 3 days post inoculation. Primers used for the PCR analysis are listed in **Table S10**.

### Construction of AT1G62914pm and AT1G62914del

To construct expression lines of *AT1G62914* with a pre-mature stop codon, we cloned the coding sequences of *AT1G62914* from *A. thaliana* Col-0 using gene-specific primers (**Table S5**). The PCR products were inserted into pENTR/D-Topo vector and the inserted gene was subsequently mutated using site-directed mutagenesis to generate *AT1G62914pm. AT1G62914pm* was cloned into the vector pEG104 using Gateway cloning (Invitrogen). To generate the *AT1G62914del* construct, two fragments of *AT1G62914* flanking the deleted region were fused at the BsaI sites and ligated into pEG104 using In-Fusion cloning.

### Stem-loop qRT-PCR

Abundance of siR1310 was quantified in *A. thaliana* or *N. benthamiana* by stem-loop qRT-PCR using 1 mg of total RNA as described in a previous study ^74^. qRT-PCR was performed using SYBR Green PCR Master Mix (Thermo Scientific) and U6 as an internal control (**Table S10**).

### Statistical analyses

Data in this study were analyzed using JMP Pro v13.0 (SAS) and presented as mean ± standard deviation (s.d.). A one-way ANOVA followed by Tukey’s HSD post hoc test was performed to compare the means of multiple groups in Fig. 4f-g, Fig. 6a-b, 6d-e and Fig. S5b. Significant differences of all pair-wise comparisons (*p* < 0.05) were indicated by different letters.

## Data availability

The raw sRNA-seq data of *A. thaliana* accession Col-0 can be found at the NCBI SRA with the accession number SRP135923. The raw sRNA-seq data of other seven *A. thaliana* accessions have been deposited in NCBI BioProject with the accession number PRJNA1037763. Source data are provided as supplementary files in this paper.

## Code availability

All the original code is available at the GitHub [GitHub Repository].

## Acknowledgements

W.M. is supported by Gatsby Charitable Foundation and the UKRI BBSRC grants BB/W00691X/1 and BBS/E/J/000PR9797. Y.H. is supported by grant from the Shanghai Collaborative Innovation Center of Agri-Seeds (ZXWH2150201). We thank Jonathan Jones, Sasha Eremina, Sophien Kamoun, and other colleagues at the Sainsbury Laboratory for critical reading of the manuscript and constructive suggestions.

## Contributions

W.M. conceived the project. L.F., A.T., Z.W., and Y.W. did the bioinformatic analyses. Y.H., B.T., H.L., and N.A did the experiments. W.M., Y.H., and L.F. analyzed the data. W.M., S.K., J.H., J.Z., and

Y.H. guided the execution of the project. W.M., Y.H., L.F., and A.T. wrote the manuscript and prepared the figures and tables. All authors commented on the paper.

## Ethics declarations

The authors declare no competing interests. A patent has been filed for using *PPR*-derived siRNAs in plant defense.

## Supplementary Figures

**Figure S1. sRNA-seq analysis of PPR-siRNAs in eight *A. thaliana* accessions.**

**a.** Size distribution of the sRNA population in each *A. thaliana* accession. **b.** Size distribution of *PPR-*siRNAs. **c.** Percentage of siRNA-producing *PPR* (TPM > 5) and non-siRNA-producing *PPR* (TPM ≤ 5) genes in Col-0 that have one or more homologs in each of the other accessions. **d.** Percentage of *PPR*-siRNAs produced from genes located in the hotspot cluster in each accession.

**Figure S2. Sequence analysis of siRNA-producing *PPRs* within the hotspot cluster in *A. thaliana*.**

**a.** *AT1G63320* contains a single intron (77 nt) that is present within the coding sequences of other siRNA-producing *PPR*s located in the hotspot cluster. Nucleotide sequence alignments show a high degree of similarity between *AT1G63320* and 11 other *PPR* genes within the hotspot region. The blue rectangle incidates exon regions and the black bar within *AT1G63320* indicates an intron. The red dashed box highlights regions in the other *PPR*s that are aligned to the intron sequence of *AT1G63320*.

**b.** Sequence logo depicting homologs of 11 sRNAs that are known to trigger *PPR*-siRNA production in *A. thaliana*.

**Figure S3. Sequence diversity analysis of siRNA-producing *PPR*s.**

**a.** Nucleotide diversity determined by Watterson’s theta and Distance in eight *A. thaliana* accessions for six gene groups: hotspot-siRNA-*PPR*s (n=14), non-hotspot-siRNA-*PPR*s (n=4), non-siRNA-*PPR*s (n=457), siRNA-*NLRs* (n=4), non-siRNA-*NLR*s (n=163), and control (n=500). “n” represents the number of genes in Col-0. Different letters label statistically significant differences. HS = hotspot. The ‘Kruskal-Wallis test’ method was used to compare the distribution of nucleotide diversity among different gene groups. The ‘Bonferroni’ method was applied for multiple-comparison correction. **b.** Distribution of Watterson’s theta and Distance values of the six gene groups. **c.** Distribution of AllenScore, presenting targeting ability of sRNA triggers, in the hotspot-siRNA-producing *PPR* and non-siRNA-*PPR* genes in each *A. thaliana* accession. A consistently lower AllenScore in the hotspot-siRNA-producing *PPR* indicates a conserved higher potential to be targeted by the trigger sRNAs. **d.** Contribution of *PPR*, *NLR* and *TAS* to the overall diversity of the 21-nt siRNA population in the eight *A. thaliana* accession.

**Figure S4. Pseudogenization of a hotspot-siRNA-producing *PPR* in the ecotype Sha.**

**a.** A region within the *PPR*-siRNA hotspot cluster in Chromosome 1 of Sha included two homologs of the *AT1G63080* in Col-0. Red, blue, and black triangles represent siRNA-producing *PPR*, non-siRNA-producing *PPR*, or non-*PPR* protein-coding genes, respectively. One of the homologs, *ATSHA-1G76750*, is predicted to encode a PPR protein, while the other, *ATSHA-1G76750-2*, is a pseudogene (presented as a yellow rectangle). Sequence alignment shows a C-to-G mutation in *ATSHA-1G76750-2* at the position 922 that leads to a premature stop codon. **b.** A limited number of 21-nt siRNAs produced from both *ATSHA-1G76750* and *ATSHA-1G76750-2*. **c.** Normalized abundance (TPM) of siRNAs that are shared or specifically produced from *ATSHA-1G76750* or *ATSHA-1G76750-2*. **d.** Phasing patterns of siRNAs produced from *ATSHA-1G76750* and *ATSHA-1G76750-2*. Arrows indicate the cleavage sites of sRNA triggers, which are conserved in the two homologs.

**Figure S5. Construction of the *62914pm* mutant line of *A. thaliana* that expresses *AT1G62914* carrying a premature stop codon.**

**a.** A single nucleotide mutation in *AT1G62914* results in a premature stop codon in the coding sequence. This mutant is named 62914pm. The blue rectangles highlight the target site of three sRNA triggers. The phasing pattern of siRNAs generated from *AT1G62914* is also presented. This mutant gene, with an N-terminal YFP tag, was introduced into a T-DNA mutant of *AT1G62914* to generate complementation lines. **b.** Relative transcript abundance of full-length *AT1G62914* in *62914mut* and two independent complementation lines (*62914pm-13* and *62914pm-15*). Values are mean ± s.d. of three biological replicates. One-way ANOVA with post hoc Tukey testing was used for statistical analysis. Different letters label statistically significant differences (*p* < 0.05). **c.** Protein products were undetectable in *62914pm-13* and *62914pm-15* plants using western blotting and an anti-GFP antibody. The black and grey arrowheads indicate putative sizes of potential YFP-AT1G62914 and YFP-62914pm protein products, respectively. The open arrowhead lables the corresponding band of 2xYFP-TurboID, which has a similar size with YFP-AT1G62914 and was used as a control.

**Figure S6. *PPR*-siRNA production is a common phenonemon in a wide range of plant species.**

**a.** siRNA-producing *PPRs* were analyzed in 55 plant species, including three early plants, ten monocots, and 42 eudicots. A phylogenetic tree was constructed using the amino acid sequence of AT2G21710. The heatmap represents number of siRNA-producing loci in each of the DYW, E, P or PLS class of PPR proteins. **b.** Genes encoding P-class PPR*s* are major sources for siRNA production in plants. Abundance of *PPR*-siRNAs spawned from genes belong to the different PPR classes. Values are average log2-tranformed TPM from all the gene members belonging to the same class.

**Figure S7. Construction of *62914del A. thaliana* abolished the production of secondary siRNAs.**

A 159 bp deletion in *AT1G62914* (*62914del*) does not affect the target sites of the sRNA triggers. The blue rectangles highlight the target sites of sRNA triggers, and the white rectangle indicates the region that is deleted in *62914del*. The region from which siR1310 is produced is highlighted by the red rectangle. The phasing pattern of siRNAs generated from wildtype *AT1G62914* transcript is also presented.

## Supplementary Tables

**Table S1.** Quantity of raw and clean reads of sRNA-seq libraries.

**Table S2.** Homologous *PPR*, *NLR*, and *TAS* genes in eight *A. thaliana* accessions.

**Table S3.** Number variations of siRNA-producing *PPRs* in eight *A. thaliana* accessions.

**Table S4.** Sequences of sRNA triggers targeting *PPR* transcripts in the eight *A. thaliana* accessions.

**Table S5.** Sequences of *AT1G62914*, *AT1G62914pm*, and *AT1G62914del*.

**Table S6.** Genome sources of plant species used in this study.

**Table S7**. 17 RefSeq proteomes used for the PPR phylogenetic analysis.

**Table S8**. PPR proteins identified in the 17 RefSeq proteomes.

**Table S9.** Analysis of siRNA-producing *PPR*s in soybean.

**Table S10**. Oligonucleotides used in this study.

## Source Data

Raw Data for results presented in Fig. 4f-g, Fig. 5b-c, Fig. 6a-b, 6d-e, and Fig. S5b.

