## Supplementary figures for "A conserved small RNA-generating gene cluster undergoes sequence diversification and contributes to plant immunity"

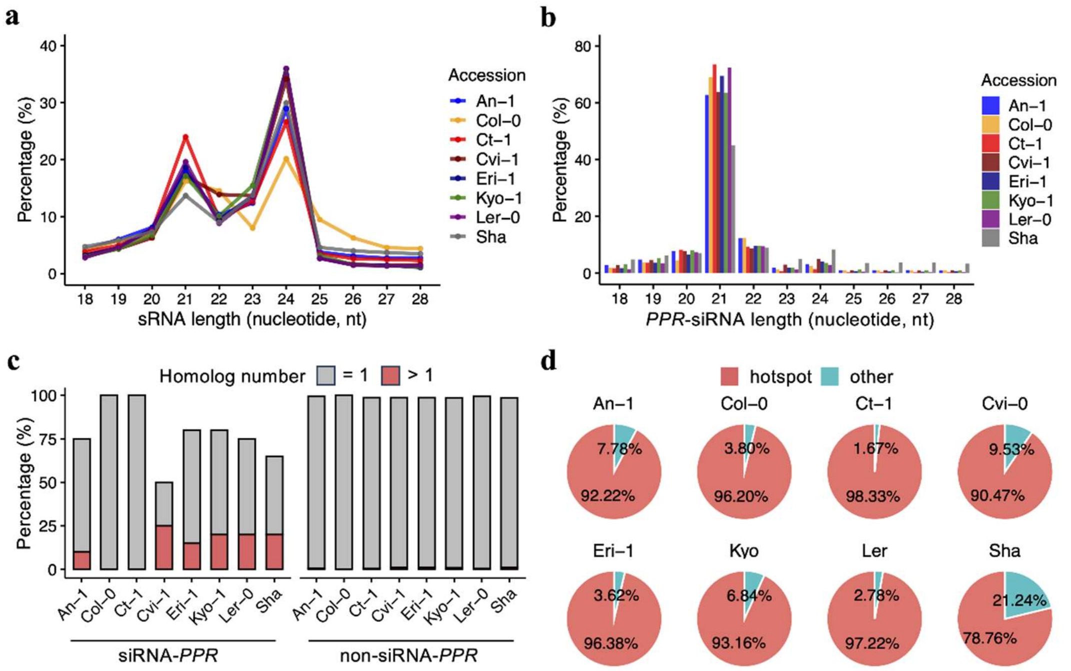


**Figure S1. sRNA-seq analysis of PPR-siRNAs in eight *A. thaliana* accessions.**

**a.** Size distribution of the sRNA population in each *A. thaliana* accession. **b.** Size distribution of *PPR*-siRNAs. **c.** Percentage of siRNA-producing *PPR* (TPM > 5) and non-siRNA-producing *PPR* (TPM ≤ 5) genes in Col-0 that have one or more homologs in each of the other accessions. **d.** Percentage of *PPR*-siRNAs produced from genes located in the hotspot cluster in each accession.

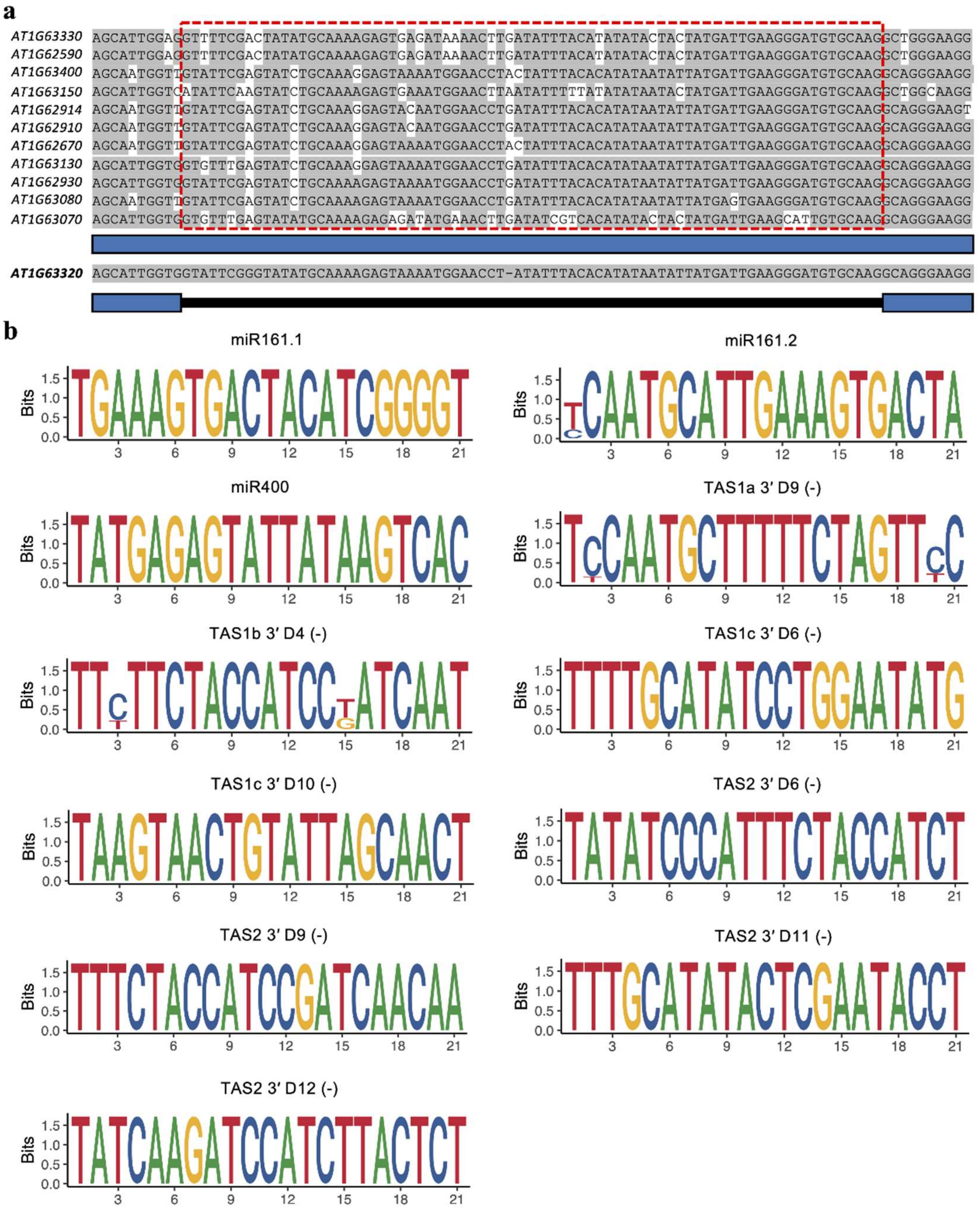


**Figure S2. Sequence analysis of siRNA-producing *PPRs* within the hotspot cluster in *A. thaliana*.**

**a.** *AT1G63320* contains a single intron (77 nt) that is present within the coding sequences of other siRNA-producing *PPR*s located in the hotspot cluster. Nucleotide sequence alignments show a high degree of similarity between *AT1G63320* and 11 other *PPR* genes within the hotspot region. The blue rectangle incidates exon regions and the black bar within *AT1G63320* indicates an intron. The red dashed box highlights regions in the other *PPR*s that are aligned to the intron sequence of *AT1G63320*. **b.** Sequence logo depicting homologs of 11 sRNAs that are known to trigger *PPR*-siRNA production in *A. thaliana*.

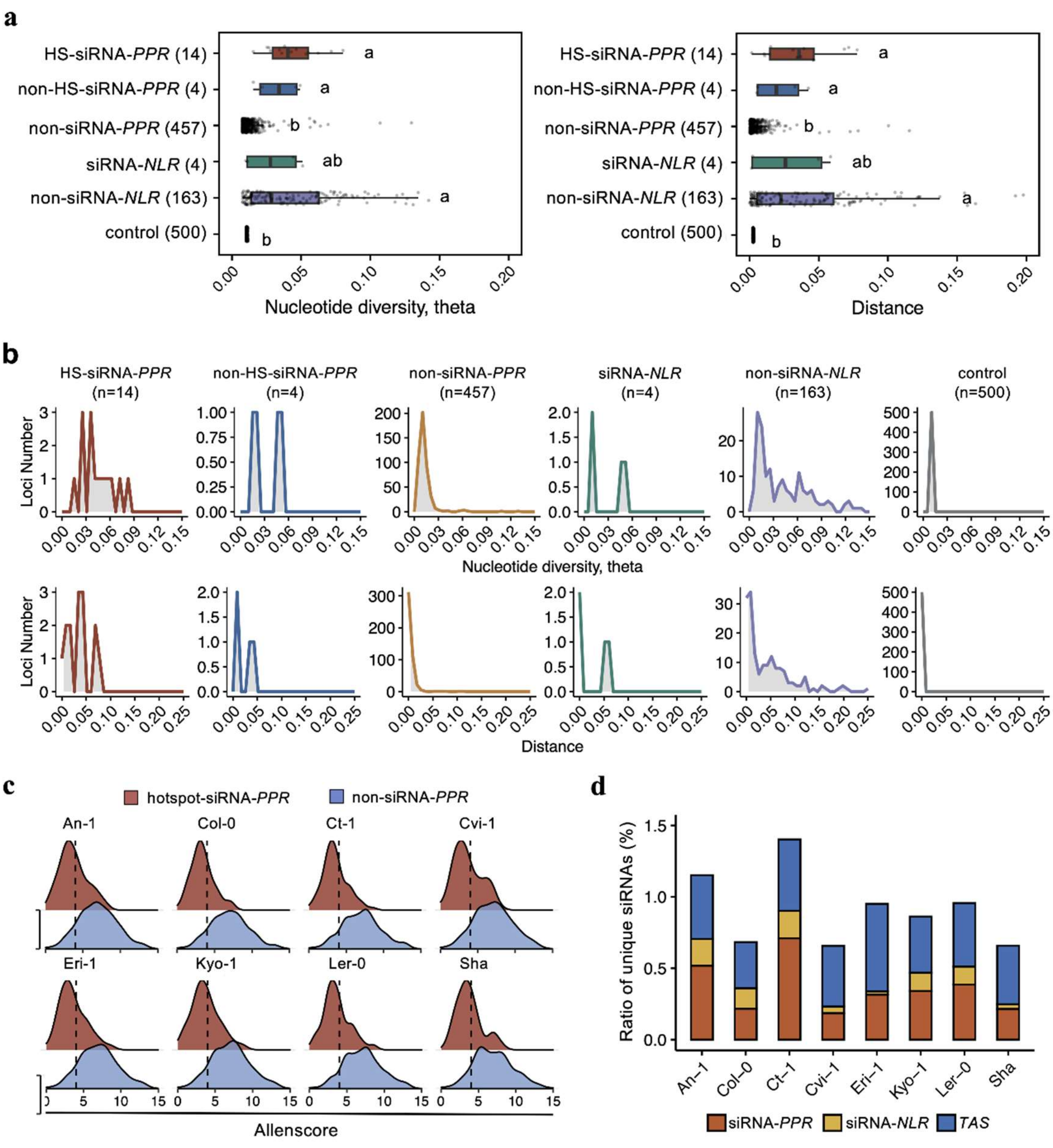


**Figure S3. Sequence diversity analysis of siRNA-producing *PPR*s.**

**a.** Nucleotide diversity determined by Watterson's theta and Distance in eight *A. thaliana* accessions for six gene groups: hotspot-siRNA-*PPR*s (n=14), non-hotspot-siRNA-*PPR*s (n=4), non-siRNA-*PPR*s (n=457), siRNA-*NLRs* (n=4), non-siRNA-*NLR*s (n=163), and control (n=500). "n" represents the number of genes in Col-0. Different letters label statistically significant differences. HS = hotspot. The 'Kruskal-Wallis test' method was used to compare the distribution of nucleotide diversity among different gene groups. The 'Bonferroni' method was applied for multiple-comparison correction. **b.** Distribution of Watterson's theta and Distance values of the six gene groups. **c.** Distribution of AllenScore, presenting targeting ability of sRNA triggers, in the hotspot-siRNA-producing *PPR* and non-siRNA-*PPR* genes in each *A. thaliana* accession. A consistently lower AllenScore in the hotspot-siRNA-producing *PPR* indicates a conserved higher potential to be targeted by the trigger sRNAs. **d.** Contribution of *PPR*, *NLR* and *TAS* to the overall diversity of the 21-nt siRNA population in the eight *A. thaliana* accession.

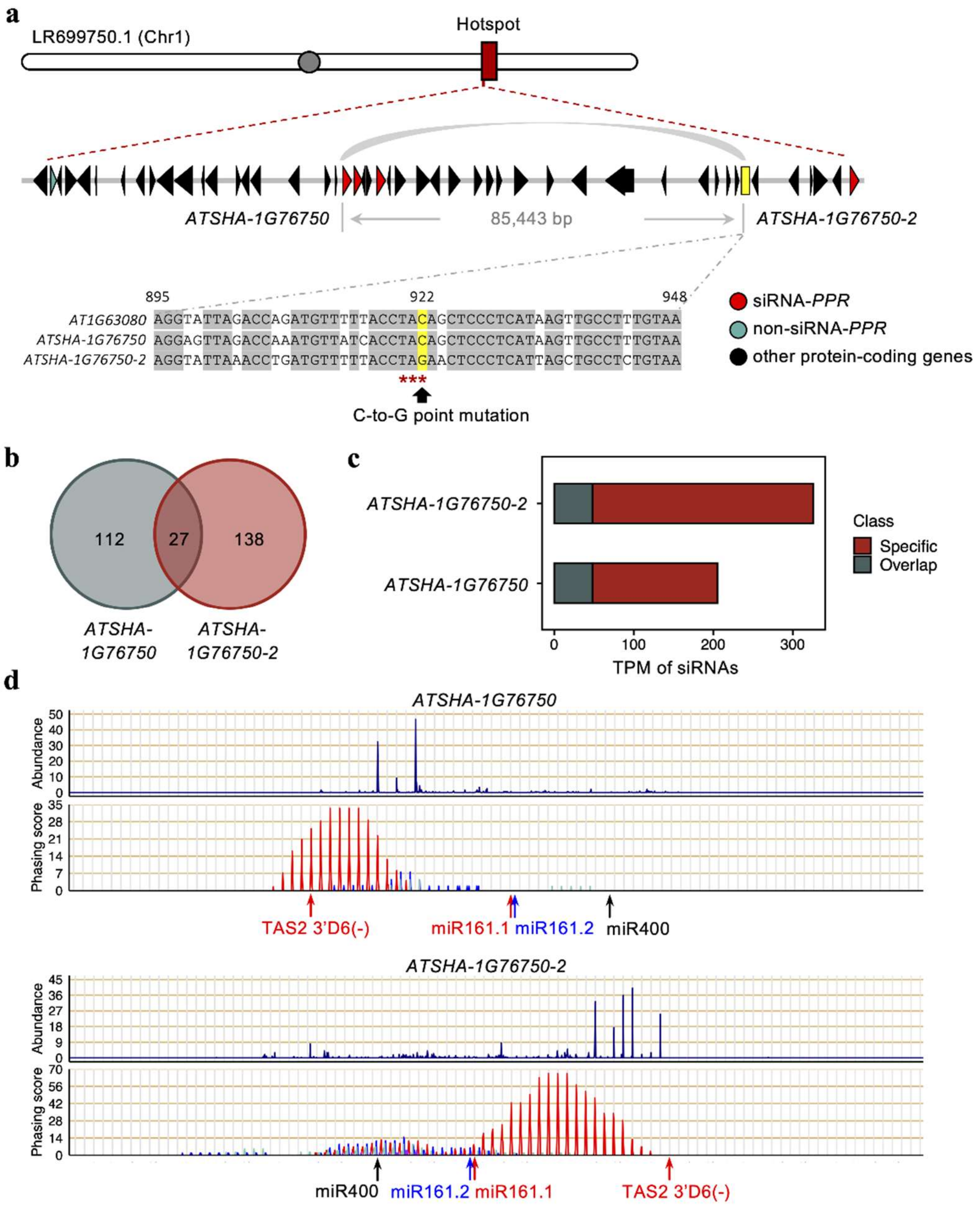


**Figure S4. Pseudogenization of a hotspot-siRNA-producing *PPR* in the ecotype Sha.**

**a.** A region within the *PPR*-siRNA hotspot cluster in Chromosome 1 of Sha included two homologs of the *AT1G63080* in Col-0. Red, blue, and black triangles represent siRNA-producing *PPR*, non-siRNA-producing *PPR*, or non-*PPR* protein-coding genes, respectively. One of the homologs, *ATSHA-1G76750*, is predicted to encode a PPR protein, while the other, *ATSHA-1G76750-2*, is a pseudogene (presented as a yellow rectangle). Sequence alignment shows a C-to-G mutation in *ATSHA-1G76750-2* at the position 922 that leads to a premature stop codon. **b.** A limited number of 21-nt siRNAs produced from both *ATSHA-1G76750* and *ATSHA-1G76750-2*. **c.** Normalized abundance (TPM) of siRNAs that are shared or specifically produced from *ATSHA-1G76750* or *ATSHA-1G76750-2*. **d.** Phasing patterns of siRNAs produced from *ATSHA-1G76750* and *ATSHA-1G76750-2*. Arrows indicate the cleavage sites of sRNA triggers, which are conserved in the two homologs.

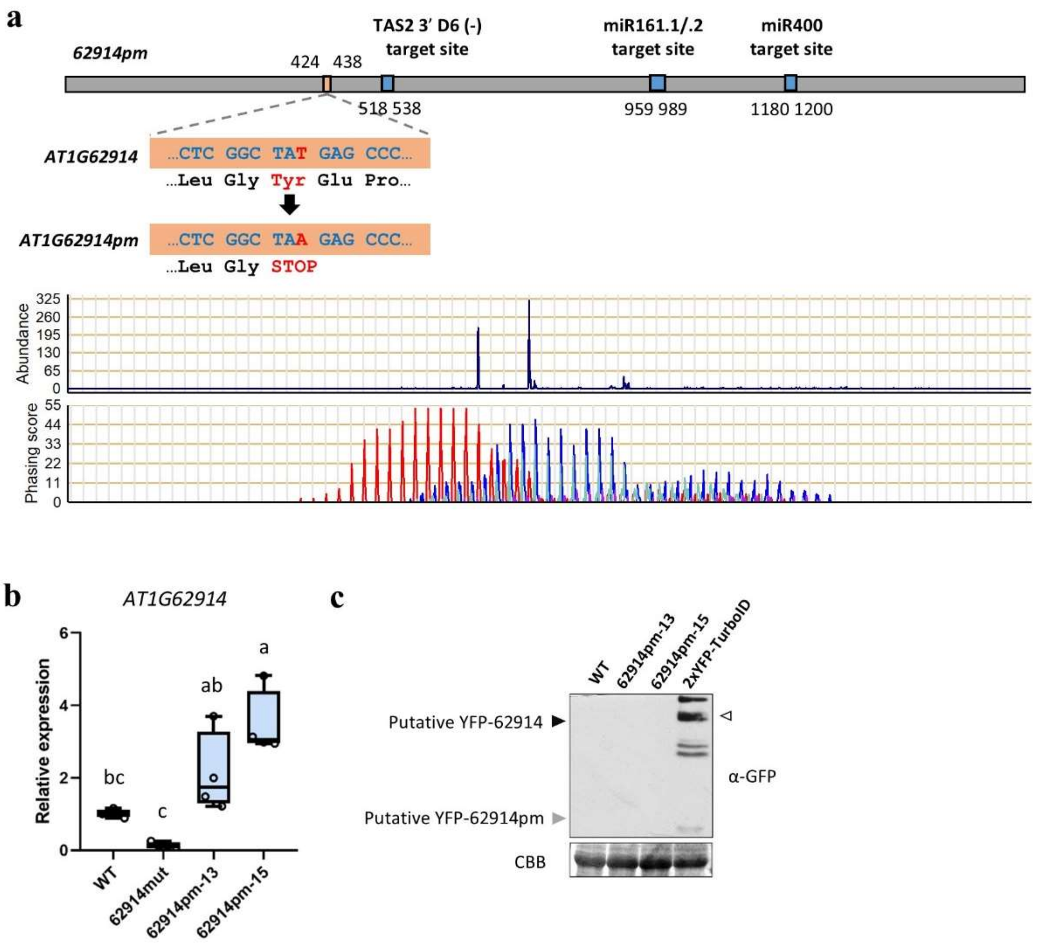


**Figure S5. Construction of the *62914pm* mutant line of *A. thaliana* that expresses *AT1G62914* carrying a premature stop codon.**

**a.** A single nucleotide mutation in *AT1G62914* results in a premature stop codon in the coding sequence. This mutant is named 62914pm. The blue rectangles highlight the target site of three sRNA triggers. The phasing pattern of siRNAs generated from *AT1G62914* is also presented. This mutant gene, with an N-terminal YFP tag, was introduced into a T-DNA mutant of *AT1G62914* to generate complementation lines. **b.** Relative transcript abundance of full-length *AT1G62914* in *62914mut* and two independent complementation lines (*62914pm-13* and *62914pm-15*). Values are mean ± s.d. of three biological replicates. One-way ANOVA with post hoc Tukey testing was used for statistical analysis. Different letters label statistically significant differences ($p < 0.05$). **c.** Protein products were undetectable in *62914pm-13* and *62914pm-15* plants using western blotting and an anti-GFP antibody. The black and grey arrowheads indicate putative sizes of potential YFP-AT1G62914 and YFP-62914pm protein products, respectively. The open arrowhead lables the corresponding band of 2xYFP-TurboID, which has a similar size with YFP-AT1G62914 and was used as a control.

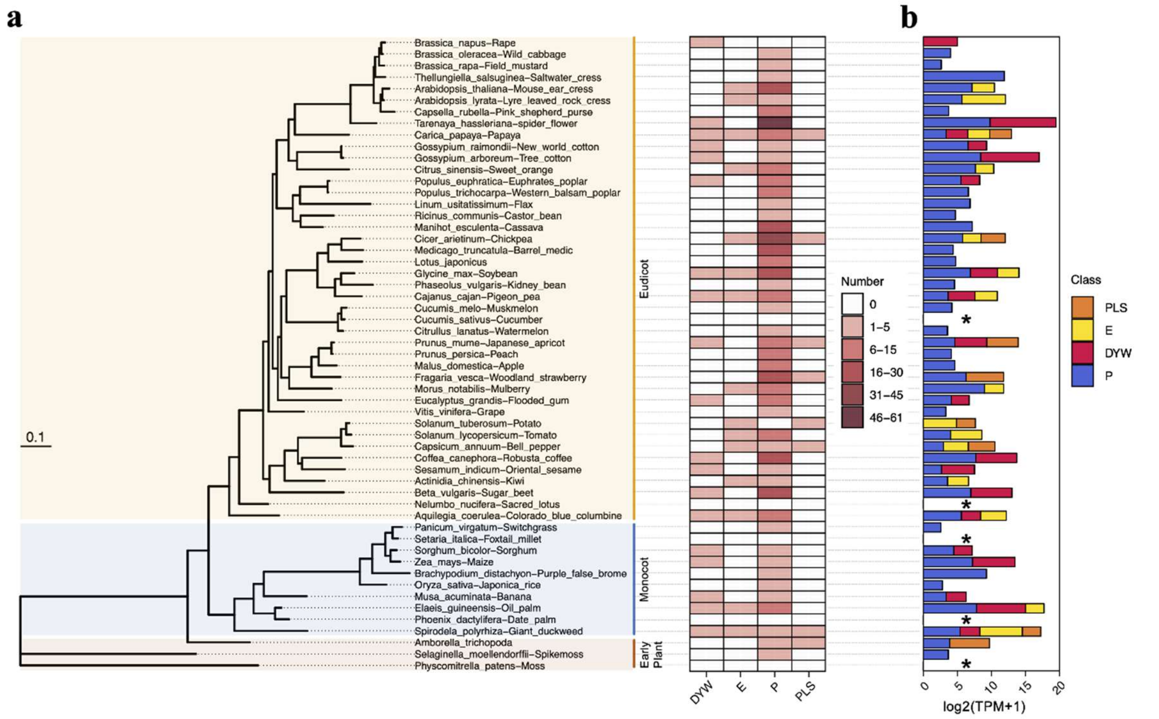


**Figure S6. *PPR*-siRNA production is a common phenonemon in a wide range of plant species.**

**a.** siRNA-producing *PPRs* were analyzed in 55 plant species, including three early plants, ten monocots, and 42 eudicots. A phylogenetic tree was constructed using the amino acid sequence of AT2G21710. The heatmap represents number of siRNA-producing loci in each of the DYW, E, P or PLS class of PPR proteins. **b.** Genes encoding P-class PPR*s* are major sources for siRNA production in plants. Abundance of *PPR*-siRNAs spawned from genes belong to the different PPR classes. Values are average log2-tranformed TPM from all the gene members belonging to the same class.

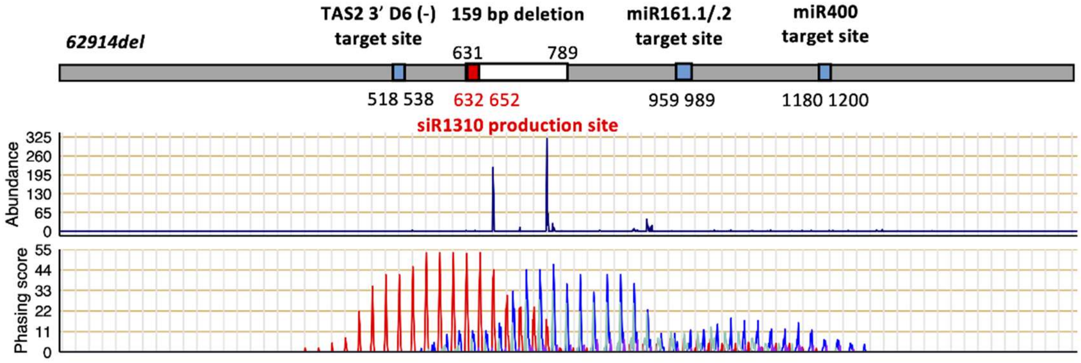


**Figure S7. Construction of *62914del A. thaliana* abolished the production of secondary siRNAs.**

A 159 bp deletion in *AT1G62914* (*62914del*) does not affect the target sites of the sRNA triggers. The blue rectangles highlight the target sites of sRNA triggers, and the white rectangle indicates the region that is deleted in *62914del*. The region from which siR1310 is produced is highlighted by the red rectangle. The phasing pattern of siRNAs generated from wildtype *AT1G62914* transcript is also presented.
